# Genome-wide conditional degron libraries for functional genomics

**DOI:** 10.1101/2024.05.29.596381

**Authors:** Eduardo Gameiro, Karla A. Juárez-Núñez, Jia Jun Fung, Susmitha Shankar, Brian Luke, Anton Khmelinskii

**Affiliations:** Institute of Molecular Biology (IMB), Mainz, Germany; Johannes Gutenberg University Mainz, Institute for Developmental Neurology (IDN), Mainz, Germany

## Abstract

The yeast knockout library, comprised of strains carrying precise deletions of individual open reading frames (ORFs), has been a vital resource to understand gene function. However, accumulation of suppressor mutations in knockout strains can lead to erroneous functional annotations. Moreover, the library is limited to non-essential ORFs, and must be complemented with hypomorphic alleles of essential ORFs for genome-wide studies. To address these limitations, we constructed genome-wide libraries of conditional alleles based on the auxin-inducible degron (AID) system for conditional degradation of AID-tagged proteins. First, we determined that N-terminal tagging is at least twice more likely to inadvertently impair protein function across the proteome. We thus constructed two genome-wide libraries with over 5600 essential and non-essential proteins fused at the C-terminus with an AID tag and an optional fluorescent protein. Almost 90% of AID-tagged proteins were degraded in the presence of the auxin analog 5-Ph-IAA, with initial protein abundance and tag accessibility as limiting factors. Genome-wide screens for DNA damage response factors with the AID libraries revealed a role for the glucose signaling factor *GSF2* in resistance to hydroxyurea, highlighting how these resources extend the toolbox for functional genomics in yeast.

## Introduction

Characterization of loss-of-function alleles is the most common approach to study gene function. With its simple genetics and broad range of resources, the budding yeast *Saccharomyces cerevisiae* is an excellent model for functional genomics (Botstein and Fink, 2011). The budding yeast knockout library was the first genome-wide collection of gene deletion strains (Giaever et al., 2002; Winzeler et al., 1999). This resource has enabled hundreds of genome-wide screens for functional profiling of the yeast genome, mapping of genetic interactions, identification of drug targets and mechanisms of drug action (Giaever and Nislow, 2014).

Despite the wide-ranging impact of the yeast knockout library, this resource has some limitations. First, because approximately 20% of yeast genes are essential (Giaever et al., 2002), i.e. their deletion is lethal, the deletion collection should be complemented with conditional alleles of essential genes for complete coverage. Libraries of essential alleles based on transcriptional repression, decreased expression by reducing gene dosage through the use of heterozygous diploids or by mRNA destabilization, temperature sensitivity or targeted protein degradation have been applied with different trade-offs, including variability in the rate and extent of downregulation or side effects of the conditions necessary for downregulation (Ben-Aroya et al., 2008; Breslow et al., 2008; Giaever et al., 2002; Kanemaki et al., 2003; Li et al., 2011; Mnaimneh et al., 2004; Snyder et al., 2019). Another limitation of the knockout library is the masking of gene-specific phenotypes by spontaneous suppressor mutations that can arise in gene deletion strains (Van Leeuwen et al, 2016; Hughes et al, 2000; Teng et al, 2013).

Here we sought to address these limitations by developing genome-wide libraries of conditional alleles based on the AID2 auxin-inducible degron system for targeted protein degradation (Nishimura et al., 2009; Yesbolatova et al., 2020). The improved AID2 system allows for tightly controlled and rapid degradation of AID-tagged proteins (Nishimura et al., 2009). We show that near complete degradation of most yeast proteins can be achieved with genome-wide AID libraries and demonstrate an application of these resources in genome-wide screens for factors involved in the DNA damage response.

## Results

### Systematic comparison of N- and C-terminal tagging

We set out to construct genome-wide auxin-inducible degron (AID) libraries, where each strain expresses a different protein fused to the AID tag. To decide at which terminus to fuse the AID tag so that the likelihood of tagging artifacts is minimized, we asked whether protein N- or C-termini are generally more amenable to tagging. Although studies of individual proteins have shown that tagging can impair protein localization or function depending on the tagged terminus, a comprehensive comparison of N- and C-terminal tagging is lacking (Weill et al., 2019).

To systematically assess the relative impact of N- and C-terminal tagging, we established a high-throughput competition assay (Fig. 1a). In this assay, pairs of strains expressing the same protein fused either at the N- or at the C-terminus to a non-fluorescent tag are co-cultured. The frequency of each cell type in the co-cultures is monitored over time by flow cytometry of strain-specific fluorescent markers, the mNeonGreen (mNG) and mCherry fluorescent proteins expressed from a strong constitutive promoter. The relative fitness (α) of the two co-cultured strains is then determined from the change in the relative frequencies of the two cell types over time (Fig. 1a, Methods). Under the assumption that tagging is either neutral or impairs fitness, but is very unlikely to improve fitness, α < 0 corresponds to reduced relative fitness of the C-terminally tagged strain and α > 0 indicates reduced relative fitness of the N-terminally tagged one.

**Figure 1.**
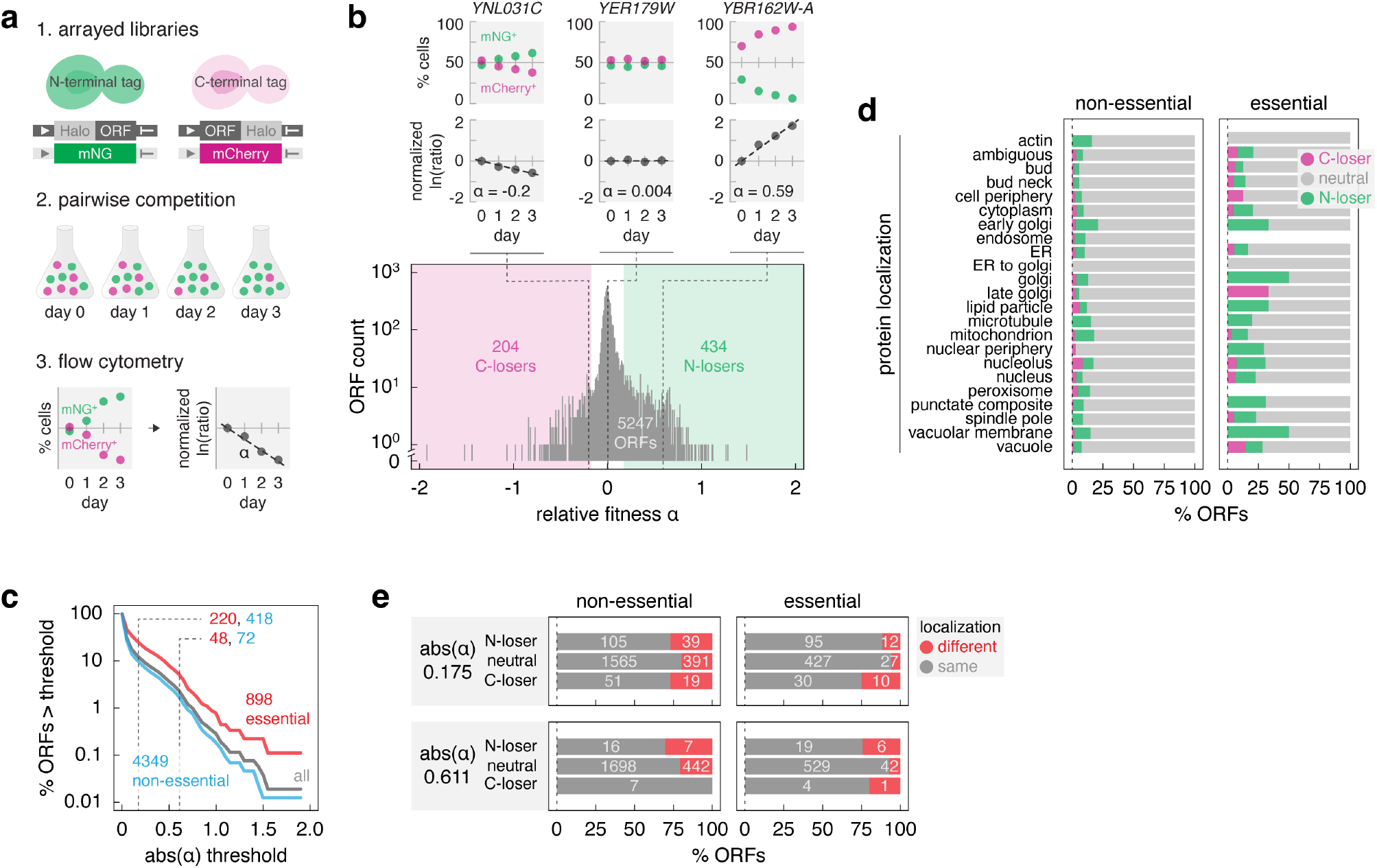
Fitness impact of N- vs C-terminal protein tagging. **a** – Competition assay to compare fitness of strains with N- or C-terminally tagged proteins. Pairs of strains with the same open reading frame (ORF) tagged at the endogenous chromosomal locus are co-cultured. The percentage of each cell type in the population is followed with flow cytometry using strain-specific fluorescent markers expressed from the constitutive *GPD* promoter (mNeonGreen (mNG), green or mCherry, magenta). The slope α of a linear fit to ln(mCherry^+^ cells/mNG^+^ cells) over time represents the relative fitness of the two strains (α > 0, competitive advantage of the ORF-Halo strain). **b** – Genome-wide assessment of N- vs C-terminal protein tagging impact on fitness. Distribution of relative fitness for 5247 ORFs assayed according to **a**. ORFs with α < -0.076 are defined as C-losers (C-terminal tagging impairs fitness, magenta area), α > 0.076 – N-losers (green area). Top, percentage of mCherry^+^ and mNG^+^ cells and linear fit to the ln(mCherry^+^/mNG^+^ cells) over time for three representative ORFs. **c** – Percentage of ORFs differentially affected by N- vs C-terminal tagging as a function of a relative fitness threshold. Number of ORFs, essential (red) or non-essential (blue), differentially affected at two thresholds (dashed lines, abs(α) = 0.175 (5% fitness difference) and 0.611 (20%), Methods) is indicated. **d** – Frequency of relative fitness differences according to protein localization. Subcellular localization based on fluorescence microscopy imaging of 3827 strains expressing C-terminally GFP-tagged proteins (Huh et al., 2003). **e** – Proteins with differential localization of N- and C-terminally GFP tagged variants (Huh et al., 2003; Weill et al., 2018), stratified by gene essentiality and differential fitness according to **b**. N-loser, neutral and C-loser ORFs defined at two differential fitness thresholds. Number of ORFs in each group is indicated.

We used the high-throughput SWAp-Tag (SWAT) approach (Meurer et al., 2018; Weill et al., 2018; Yofe et al., 2016) to tag open reading frames (ORFs) at endogenous chromosomal loci with a sequence encoding the Halo tag (Los et al., 2008). The SWAT approach makes use of strain arrays with individual ORFs marked with an acceptor module. This module is inserted immediately before the stop codon in C-SWAT strains or after the start codon in N-SWAT strains, except for ORFs encoding proteins with predicted N-terminal signal peptides or N-terminal mitochondrial targeting signals where the acceptor module is placed after the predicted signal sequences. Using semi-automated crossing, the acceptor module can be replaced with practically any tag or regulatory element provided on a donor plasmid (Yofe et al., 2016). We chose the Halo tag due its size (33 kDa), similar to many commonly used fluorescent protein tags and to the mNG-AID*-3myc tag in the AID- v1 library described below, and lack of evidence for a dominant negative effect on the tagged proteins. Halo tagging using N-SWAT and C-SWAT libraries was efficient, yielding homogeneous strains with over 99% of the cells in most strains expressing the expected fusion protein under control of its native regulatory elements (Fig. S1a, Methods). As the resulting Halo-ORF and ORF-Halo libraries are haploid, the ORF tagged in each strain is the only source of that specific protein.

Next, we competed pairs of matched N- and C-terminally tagged strains and determined the relative fitness impact of the Halo tag location (Fig. 1a). The competition assay yielded reproducible estimates of relative fitness (α) (Fig. S1b). The distribution of relative fitness across all assayed pairs was centered at 0.02 ± 0.17 (mean ± s.d., n = 5247 ORFs) (Fig. 1b, Table S1), suggesting that the impact of the Halo tag was generally independent of the tagged terminus. Importantly, the strain-specific fluorescent markers affected fitness to a similar degree, as indicated by the relative fitness of 0.05 ± 0.01 (mean ± s.d., n = 92 technical replicates) for a pair of untagged strains expressing mNG or mCherry (Fig. S1c). Therefore, the contribution of the fluorescent makers to the detected fitness differences is likely negligible. Based on these observations, we estimate that fitness differences as low as 5% (corresponding to an absolute α = 0.175, Methods) can be measured in the competition assay. Hereafter, ORFs with α > 0.175 are referred to as N-losers, with α < -0.175 as C-losers and with absolute α ≤ 0.175 as neutral.

Tagging of non-essential ORFs, especially those whose deletion has little to no impact on growth, is unlikely to impair fitness. Accordingly, essential ORFs were more likely to exhibit differential fitness defects in the competition assay (24% compared to 10% of non-essential ORFs) (Fig. 1c). This suggests that up to a quarter of the proteome is differentially affected by terminal tagging, under the assumption that tagging similarly affects essential and non-essential proteins. In addition, tagging at both termini can be detrimental to the function of some proteins. Such cases are likely rare, as no single instance could be detected in triple competition assays for a random set of 45 ORFs, where both tagged variants were competed against the wild type (Fig. S1d, e).

Whereas 155 out of 898 essential ORFs were N-losers, exhibiting reduced relative fitness with N-terminal compared to C-terminal tagging, only 65 essential ORFs were C-losers. This tendency held across the whole dataset, with a total of 434 N-losers and 204 C-losers (Fig. 1b), indicating that N-terminal tagging is more likely to impair protein function compared to C-terminal tagging. Interestingly, the frequency of differential fitness defects varied between subcellular compartments (Fig. 1d). Proteins of the golgi, vacuolar membrane and lipid particle were most likely to be differentially affected by terminal tagging, suggesting that terminal tagging could affect protein localization. Supporting this notion, differential fitness defects correlated with differential localization of N- and C-terminally tagged proteins (Fig. 1e). Using datasets of subcellular localizations for most yeast proteins determined with fluorescence microscopy of strains expressing N- and C-terminal GFP fusions (Breker et al., 2013; Huh et al., 2003; Weill et al., 2018), we observed that for essential genes, only 6% of neutral ORFs exhibited different localizations of N- and C-terminally tagged variants, whereas this fraction increased to 11% and 25% for N-losers and C-losers, respectively (Fig. 1e). The magnitude of this trend is likely underestimated considering that for 636 proteins the N-terminal Halo tag was not placed at the very N-terminus but after the predicted N-terminal signal peptide or mitochondrial targeting signal, to avoid blocking these localization signals and increase the likelihood of correct protein localization (Yofe et al., 2016). Optimized tagging of proteins with these N-terminal localization signals likely also contributes to the lack of correlation between differential fitness defects and occurrence of terminal localization signals (Fig. S1f, Table S2). Taken together, we conclude that while N-terminal tagging can be beneficial in individual cases, C-terminal tagging is overall less likely to affect protein localization and function. Nevertheless, further work is needed to understand how the type of tag, its size and biophysical properties, and the linker between the tag and the protein of interest affect protein localization and function across the proteome.

### AID libraries

Based on the above analysis, we constructed genome-wide C-terminally tagged AID libraries. We combined the short AID* tag, which consists of amino acids 71-114 from AtIAA17 (Morawska and Ulrich, 2013; Nishimura et al., 2009), with the AID2 system (Yesbolatova et al., 2020) (Fig. 2a). The AID2 system relies on the OsTir1 F-box protein with the F74G mutation and a ligand, the auxin analog 5-phenyl-indole-3-acetic acid (5-Ph-IAA), to achieve quick and efficient degradation of AID-tagged proteins at significantly lower ligand concentrations than the original AID system and without basal degradation in the absence of 5-Ph-IAA (Nishimura et al., 2009; Yesbolatova et al., 2020).

**Figure 2.**
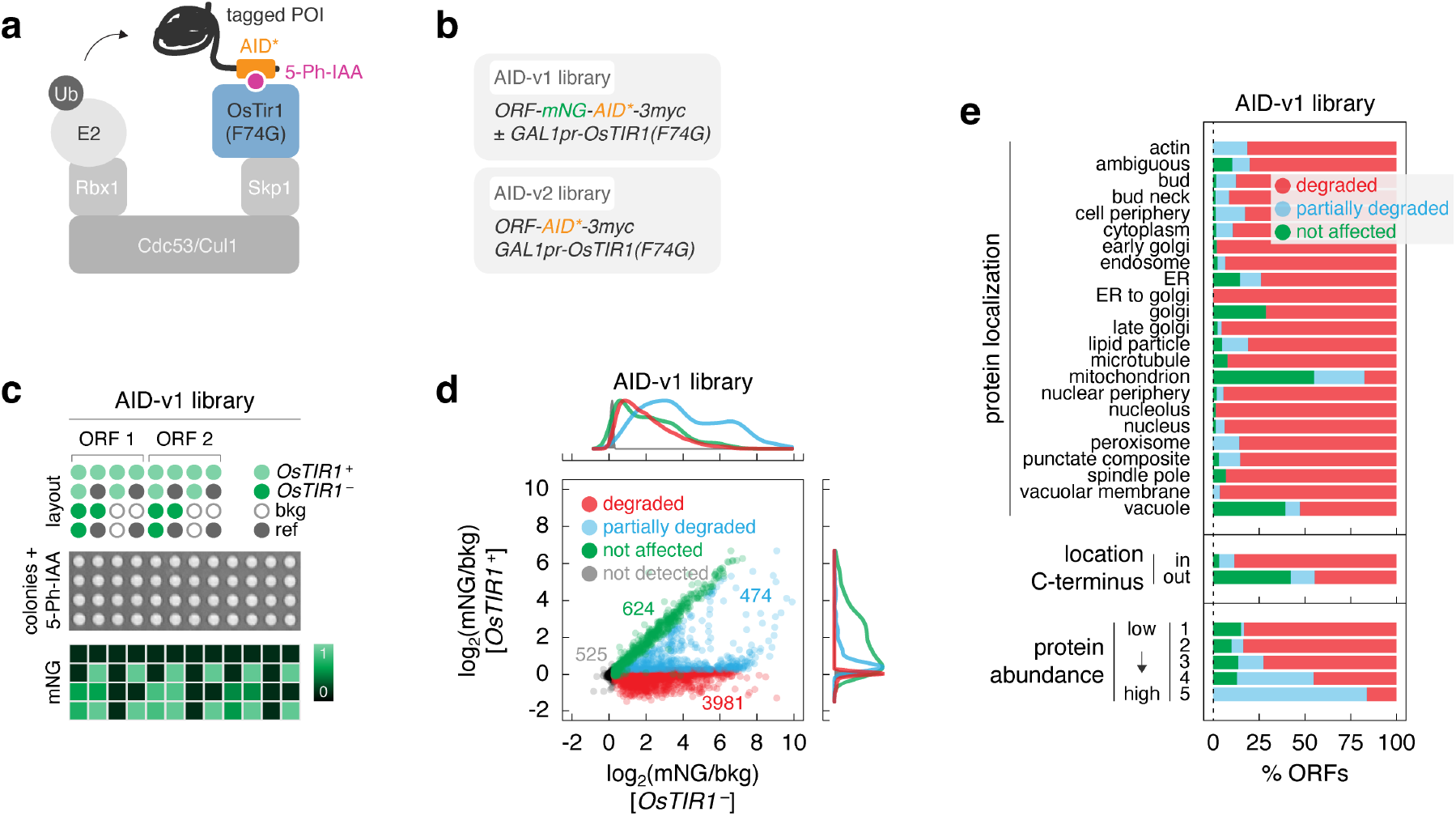
Efficient protein degradation with genome-wide AID libraries. **a** – Features of the auxin-inducible degron system in the AID libraries. A protein of interest (POI) with a C- terminal AID* tag can be ubiquitinated and degraded in the presence of 5-Ph-IAA upon expression of the F- box protein OsTir1. **b** – Different tags used in the AID libraries. Expression of *OsTIR1* is controlled by the galactose-inducible *GAL1* promoter. **c** – Characterization of the AID-v1 library with fluorescence measurements of colony arrays. Top, for each ORF, mNG-AID*-3myc-tagged strains with (+) or without (-) the *GAL1pr-OsTIR1* construct were placed next to each other. A reference strain (ref, with an abundant mNG-tagged protein, for correction of spatial and plate effects) and a strain without a fluorescent tag (bkg, for background correction) were included in each 4x4 group. Middle, colony arrays were grown on galactose medium with 1 μM 5-Ph-IAA. Bottom, example heatmap of mNG fluorescence intensities. **d** – Protein levels (mNG signal in units of background fluorescence, mNG/bkg) in the AID-v1 library determined according to **c**. Not detected, strains without a detectable mNG signal in the absence of *OsTIR1* (mNG/bkg(*OsTIR1*^-^) ≤ 1.2). Detectable proteins were classified by the extent of *OsTIR1*-dependent degradation (Methods). Top and right, normalized frequency distributions of protein levels in each category. **e** – Frequency of *OsTIR1*-dependent degradation phenotypes in the AID-v1 library according to protein localization, location of the C-terminus (cytosol – in, lumen of organelles or extracellular – out) for membrane proteins according to (Kim et al., 2006) or abundance of the mNG-AID*-3myc-tagged proteins.

Expression of OsTir1(F74G) from the strong constitutive *GPD* promoter resulted in an obvious fitness defect, whereas expression from the weaker *CYC1* and *ADH1* promoters did not affect fitness and conditional expression from the strong *GAL1* promoter had a minor impact on fitness (Fig. S2a). We decided to use the *GAL1-OsTIR1(F74G)* construct for the AID libraries as the conditional nature of the *GAL1* promoter is likely to limit adaptation to expression of OsTir1(F74G) and the high expression levels of OsTir1(F74G) are less likely to limit degradation of AID-tagged proteins. Using C-SWAT, we tagged each ORF with mNG-AID*-3myc (AID- v1 library, 5604 tagged ORFs) or with AID*-3myc (AID-v2 library, 5620 tagged ORFs) and concomitantly introduced the *GAL1pr-OsTIR1(F74G)* expression construct (Fig. 2b, S2b). The 3myc epitope tag can be used to determine protein levels with immunoblotting, whereas the mNG tag in the AID-v1 library allows assessing protein levels with fluorescence measurements. Whereas all strains in the AID-v2 library carry the *OsTIR1* construct, the AID-v1 library comprises two strains for each ORF, with or without *OsTIR1* (*OsTIR1*^+^ and *OsTIR1*^-^, respectively) (Fig. 2b).

We assessed the frequency and the extent of AID-dependent degradation across the proteome using mNG fluorescence in the AID-v1 library. Strains were grown as ordered colony arrays on agar plates with galactose and 1 μM 5-Ph-IAA, and their fluorescence was measured with a plate reader to determine the levels of mNG- AID*-3myc-tagged proteins (Fig. 2c, Methods). We estimated the detection limit of the colony fluorescence assay at approximately 200 molecules per cell (Fig. S2c). Protein expression levels in *OsTIR1*^-^ strains correlated well with independent estimates of protein abundance (Meurer et al., 2018) (Fig. S2d), indicating that the AID* tag has little impact on protein levels in the absence of the cognate F-box protein. Out of 5079 proteins detected in *OsTIR1*^-^ strains, 4455 (∼88%) were significantly depleted in *OsTIR1*^+^ strains (Fig. 2d, Table S3). 3981 proteins could not be detected specifically in the *OsTIR1*^+^ background. Hereafter, we will refer to these proteins as degraded, although it is likely that at least in some cases degradation is not complete but the remainder is below the detection limit of our plate reader assay. Nevertheless, 474 proteins were unequivocally degraded only partially, as they were detectable in the *OsTIR1*^+^ background but at reduced levels compared to the *OsTIR1*^-^ background (Fig. 2d). We verified this classification by immunoblotting in time-course experiments. Proteins from the degraded group were depleted to below the detection limit within 60 min of 5-Ph-IAA addition, partially degraded proteins were depleted less or exhibited a degradation-resistant pool, and levels of not affected proteins remained stable over time (Fig. S2e). The fraction of conditionally degraded proteins (degraded or partially degraded) varied with subcellular localization, and was lowest for proteins localized to membrane-bound organelles such as mitochondria, vacuole, golgi and endoplasmic reticulum (ER) (Fig. 2e), likely due to inaccessibility of the AID* tag to the cytosolic OsTir1. Supporting this notion, membrane proteins with cytosolic C-termini were more likely to be conditionally degraded compared to those with lumenal or extracellular C-termini (Kim et al., 2006) (Fig. 2e). Finally, the extent of AID-dependent degradation varied with initial protein abundance, in that highly expressed proteins were more likely to be only partially degraded compared to lowly expressed ones (Fig. 2e, Fig. S2f), suggesting saturation of the AID machinery. It remains to be determined whether OsTir1 or the yeast subunits of the SCF ubiquitin ligase are limiting factors for degradation of abundant proteins.

We asked whether AID-dependent protein degradation results in cellular phenotypes. For that, we measured colony sizes in the arrayed AID-v1 library as a readout of strain fitness (Fig. 2c, Methods). We first quantified the fitness impact of OsTir1 expression by comparing colony sizes of *OsTIR1*^-^ and *OsTIR1*^+^ strains for non-essential ORFs. As degradation of non-essential proteins is not expected to affect fitness, the difference in colony size between *OsTIR1*^-^ and *OsTIR1*^+^ backgrounds can be attributed to OsTir1 expression. On average, the presence of the *OsTIR1* construct reduced colony size by 7% (Fig. S3a). The same difference was observed for the 624 ORFs in the not affected group, where AID-tagged proteins were not degraded in the *OsTIR1*^+^ background. To determine how AID-dependent protein degradation affects fitness, we then corrected colony sizes of *OsTIR1*^+^ strains by the fitness impact the *OsTIR1* construct (Methods). After the correction, we found that ∼12% of *OsTIR1*^+^ strains (675 out of 5604) exhibited impaired fitness compared to *OsTIR1*^-^ strains (Fig. 3a, Table S3). As expected, the frequency of fitness defects was higher for essential proteins. Whereas only ∼4% of *OsTIR1*^+^ strains for non-essential ORFs (176 out of 4698) had impaired fitness, fitness defects were observed for over 55% of essential ORFs (499 out of 906) (Fig. 3b). A similar frequency was previously observed with a set of AID alleles constructed for 758 essential ORFs using the original AID system (Snyder et al., 2019). However, over a third of these alleles exhibited fitness defects even in the absence of auxin, which were further compounded by off-target effects of auxin.

**Figure 3.**
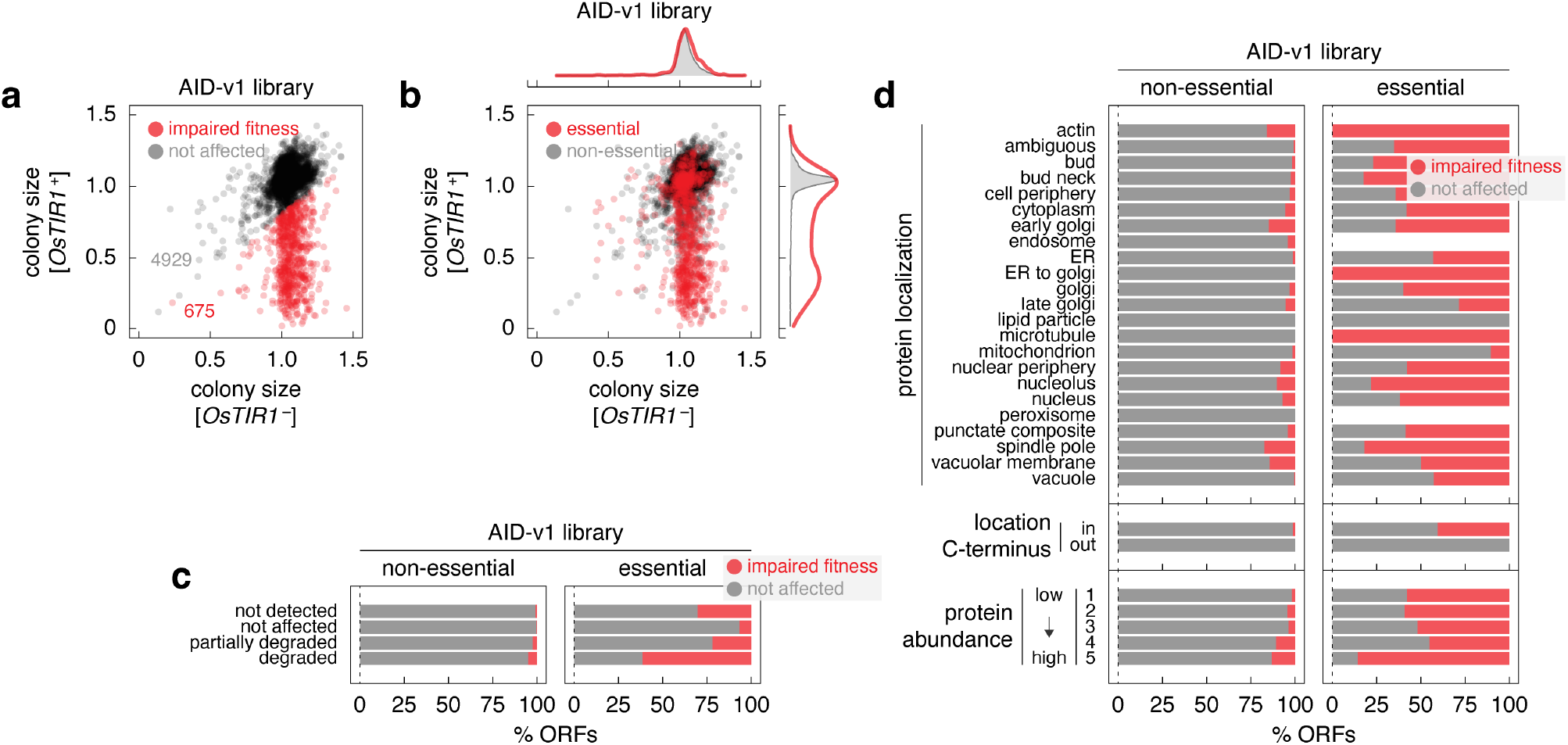
Fitness defects in the AID libraries upon induction of protein degradation. **a, b** – Normalized colony sizes in the AID-v1 library grown according to Fig. 2c. **b**, essential and non-essential ORFs are marked; top and right, normalized frequency distributions of colony sizes in each category. **c, d** – Frequency of *OsTIR1*-dependent fitness defects in the AID-v1 library according to the extent of *OsTIR1*- dependent degradation phenotypes (**c**) or according to localization, location of the C-terminus (cytosol – in, lumen of organelles or extracellular – out) or abundance of the mNG-AID*-3myc-tagged proteins (**d**).

Importantly, the frequency of fitness defects in the AID-v1 library correlated with the extent of protein degradation. Whereas degradation resulted in a fitness defect for over 61% of essential proteins, this fraction dropped to 22% and 7% of essential proteins that were partially degraded or not degraded in an AID-dependent manner, respectively (Fig. 3c). Interestingly, degradation of 38% of essential proteins did not result in a fitness defect. We considered three explanations for this observation. First, it is possible that in some cases degradation is incomplete, resulting in low protein levels that are below the detection limit of our assay but are sufficient for viability. As proteins normally expressed at low levels are more likely to fall below the detection limit even when partially depleted, we asked whether strains with essential and degraded proteins but without a fitness defect are enriched in low-abundance proteins. However, there was no difference in protein levels between *OsTIR1*^-^ strains for essential and degraded proteins with and without a fitness defect in the *OsTIR1*^+^ background (Fig. S3b). Second, we considered the nature of the essential genes in these two groups. Namely, we compared the frequency of core essential genes, which are always required for viability, and conditional essential genes, which vary in essentiality depending on the genetic background or environment (Bosch-Guiteras and van Leeuwen, 2022). Interestingly, the set of essential and degraded proteins without an accompanying fitness defect was enriched in conditional essential genes defined by two independent measures: essentiality across *S. cerevisiae* natural isolates (Peter et al., 2018) or with bypass suppression interactions in a laboratory strain (van Leeuwen et al., 2020) (Fig. S3c, odds ratio = 1.6, p-value = 0.04 in a Fisher’s exact test and odds ratio = 1.7, p-value = 0.02, respectively). This suggests that conditional essentiality could explain the observed lack of fitness defects upon degradation of some essential proteins. Third, it is likely that the fraction of strains with the degradation phenotype is overestimated in our analysis as colony fluorescence usually scales with colony size, which would result in underestimated fluorescence levels for strains with fitness defects (Fung et al., 2022). Indeed, *OsTIR1*^+^ strains with impaired fitness typically exhibited fluorescence levels below background autofluorescence (Fig. S3d, e). Finally, the frequency of fitness defects depended on the accessibility of the tag and the initial abundance of the tagged proteins (Fig. 3d), in agreement with the analysis of the degradation extent (Fig. 2e). Thus, while the likelihood and the extent of conditional protein degradation in the AID libraries depend on the abundance of the target protein and the accessibility of the AID tag, most yeast proteins can be efficiently degraded with the AID system, resulting in cellular phenotypes consistent with loss of the target protein.

### Screens for DNA damage response factors

We sought to apply the AID libraries to screen for factors involved in the DNA damage response using sensitivity to genotoxic agents as a phenotypic readout. We decided to use the DNA damaging agent methyl methanesulfonate (MMS), the topoisomerase I inhibitor camptothecin (CPT) and the ribonucleotide reductase inhibitor hydroxyurea (HU) as genotoxic agents and the AID-v2 library, reasoning that an overall smaller tag (without the mNG moiety) is less likely to inadvertently affect protein function. The library was grown as ordered colony arrays on galactose media with 5-Ph-IAA, a genotoxic agent or both compounds (Fig. 4a). First, we observed that treatment of the AID-v2 library with 1 μM 5-Ph-IAA resulted in fitness defects consistent with the AID-v1 library (spearman correlation coefficient ρ = 0.6, Fig. S4a). These fitness defects similarly correlated with protein essentiality, expression levels, tag accessibility and extent of AID-dependent degradation (Fig. S4b-e), indicating that the AID-v1 and AID-v2 libraries perform comparably.

**Figure 4.**
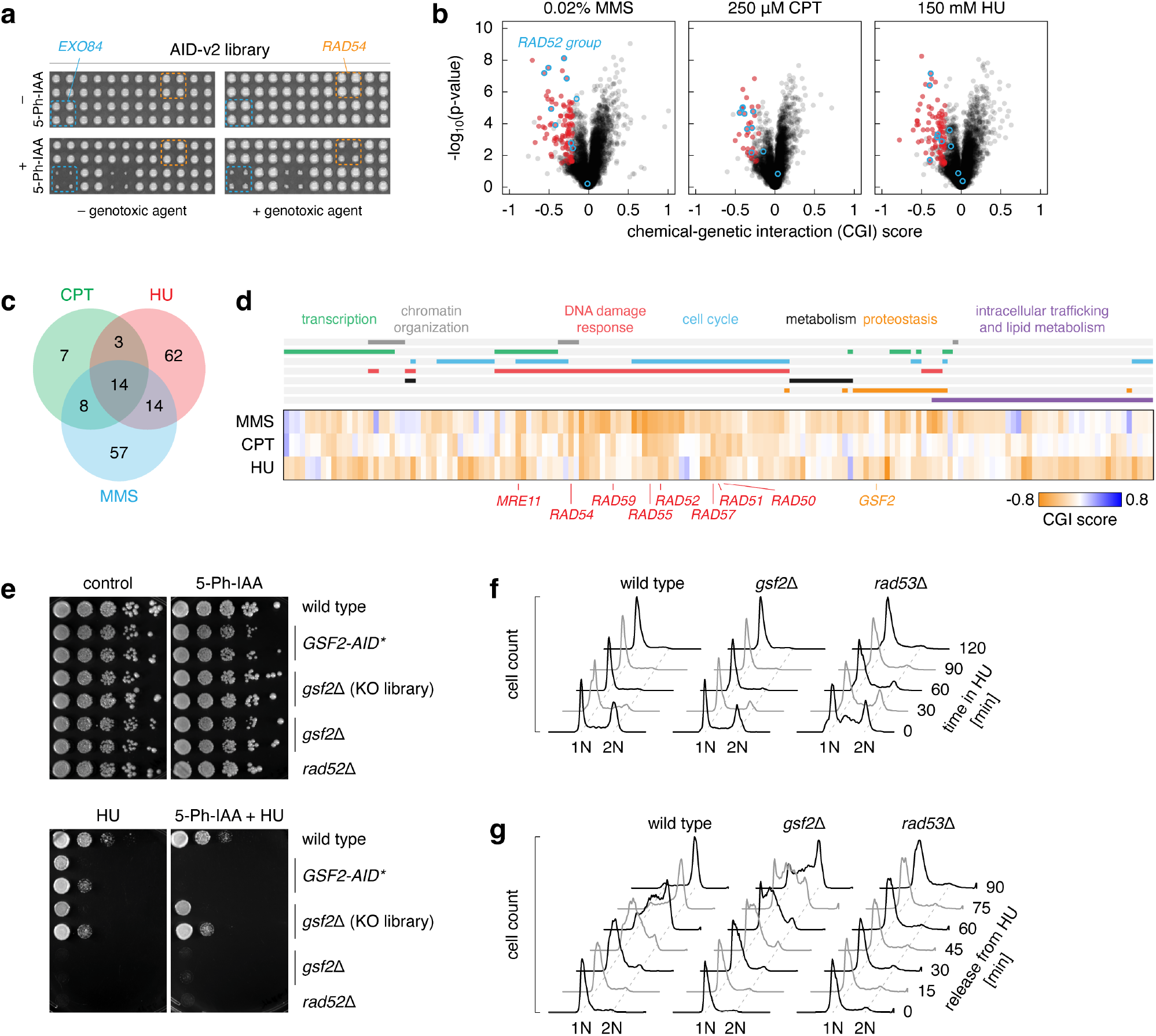
Genome-wide screens for DNA damage response factors. **a** – Arrayed fitness screens. The AID-v2 library was grown on galactose media with or without MMS, CPT or HU as genotoxic agents (GAs) and 1 μM 5-Ph-IAA. **b** – Volcano plots of chemical-genetic interaction (CGI) scores between the AID alleles and the indicated GAs. Negative chemical-genetic interactors with each genotoxic agent (red, colony size (5-Ph-IAA + GA) ≤ 0.9, CGI score ≤ 0.2 and p-value < 0.05) and *RAD52* epistasis group genes (blue) are marked. **c, d** – Venn diagram (**c**) and heatmap (**d**) of negative chemical-genetic interactors (red in **b**) with the three genotoxic agents. **d**, functional categories based on gene ontology terms (top, Methods) and *RAD52* epistasis group genes (bottom) are indicated. **e** – Sensitivity of *GSF2* mutants to HU. Ten-fold serial dilutions of the indicated strains (GSF2-AID* strain from the AID-v2 library, *gsf2*Δ mutant from the haploid yeast knockout library and an independently constructed *gsf2*Δ strain) on galactose media with 5-Ph-IAA, HU or both compounds. **f, g** – Flow cytometry analysis of DNA content. Asynchronous cultures grown in galactose medium were treated with HU at time point zero (**f**). HU-arrested cultures were released into galactose medium without HU (**g**). G1 and G2 peaks are labeled 1N and 2N, respectively.

Next, we calculated chemical-genetic interaction (CGI) scores between each AID allele and a genotoxic agent (Fig. S5a). We focused on negative CGIs whereby the double perturbation (5-Ph-IAA and genotoxic agent) results in a stronger fitness defect than expected from the combined effects of the single perturbations (Methods) (Baryshnikova et al., 2010b; Parsons et al., 2003). This led to the identification of 93, 32 and 93 potential resistance factors (CGI score ≤ -0.2 and p-value < 0.05) for MMS, CPT and HU, respectively (Fig. 4b, Table S4). There was substantial overlap between the non-essential factors identified here and previous screens with the yeast knockout library (Fig. S5b) (Chang et al., 2002; Parsons et al., 2003). Nevertheless, AID alleles of most resistance factors identified with knockout strains were not sensitive to the corresponding genotoxic agents. We asked whether these factors might not be degraded in the respective AID strains. However, the frequency of ORFs with AID-dependent degradation was similar for screen-specific and shared resistance factors (Fig. S5b). Differences in growth conditions and definitions of significant fitness defects likely account for the screen-specific CGIs. There were 58 essential genes among the combined 165 potential resistance factors identified in the three screens (Fig. S5c). CGIs for both essential and non-essential genes were reproducible in spot tests (Fig. S5d). Additional essential factors could in principle be identified in further screens with reduced 5-Ph-IAA concentrations, which allow determining CGIs for strains that otherwise exhibit no growth in 1 μM 5-Ph-IAA (Fig. S5e). Taken together, these results validate the screening approach with AID libraries.

Although the three genotoxic agents act through distinct mechanisms, all can result in the accumulation of DNA double strand breaks (DSBs) (Li et al., 2017; Singh and Xu, 2016; Wyatt and Pittman, 2006). Accordingly, shared resistance factors for MMS, CPT and HU included most members of the *RAD52* epistasis group (Fig. 4b, c), which are involved in repair of DSBs (Game and Mortimer, 1974; Resnick, 1969; Symington, 2002). Moreover, most of the identified resistance factors were associated with gene ontology terms related to DNA damage response, transcription, chromatin organization and cell cycle (Fig. 4d, Fig. S5c, Methods). Notably, whereas CPT- and MMS-specific factors were enriched in genes associated with cell cycle progression, DNA damage response and transcription, HU-specific factors were enriched in components of the endomembrane system, including proteins of the endoplasmic reticulum (ER) and components of endosomal sorting complexes required for transport (ESCRT) (Fig. 4d, Fig. S5c, Table S5). This suggests that HU impairs proliferation through multiple mechanisms, consistent with prior reports (Huang et al., 2016; Kuong and Kuzminov, 2009; Musiałek and Rybaczek, 2021; Takano and Nakatsukasa, 2023).

Finally, we examined *GSF2*, one of the HU-specific resistance factors identified in the screen (Fig. 4d, Fig. S5c). Gsf2 is an integral membrane protein localized to the ER that is involved in biogenesis and export of the hexose transporter Hxt1 from the ER (Sherwood and Carlson, 1999). We confirmed that the *GSF2-AID\** strain from the AID-v2 library is sensitive to HU specifically in the presence of 5-Ph-IAA (Fig. 4e). A *gsf2*Δ mutant from the haploid yeast knockout library (Giaever et al., 2002; Winzeler et al., 1999) was not sensitive to HU. In contrast, a *gsf2*Δ strain generated by sporulation of a heterozygous diploid knockout exhibited HU sensitivity similar to the *GSF2-AID\** strain (Fig. 4e). This suggests that additional mutations acquired by the knockout library strain are masking the HU sensitivity phenotype. To gain insights into the role of *GSF2* in resistance to HU, we analyzed how HU affects cell cycle progression of the *gsf2*Δ mutant. Knockout of *GSF2* did not prevent yeast cells from arresting in early S-phase in response to HU (Fig. 4f), suggesting that *GSF2* plays no role in the activation of the DNA replication checkpoint. However, the *gsf2*Δ mutant exhibited delayed cell cycle progression after release from HU (Fig. 4g). This phenotype was distinct from a strain lacking the checkpoint kinase Rad53, which is essential for recovery from DNA replicational stress (Bell and Labib, 2016; Desany et al., 1998; Tercero and Diffley, 2001). These observations point towards a role of *GSF2* in recovery from the checkpoint arrest but the underlying mechanism remains to be determined.

In conclusion, our work establishes genome-wide libraries of conditional auxin-inducible degron alleles in budding yeast. We demonstrate how these resources can be applied to uncover gene functions, thus adding to the toolkit for functional genomics in this organism (Botstein and Fink, 2011). While the extent of protein degradation in the AID libraries can be assessed in a high-throughput manner with mNG fluorescence, the optional nature of the mNG moiety expands the scope of potential applications. The AID-v2 library should be ideal for high-content fluorescence microscopy screens, where the mNG moiety could otherwise limit screen design. A similar resource was independently constructed and characterized by Valenti et al. (Valenti et al., 2024). In the future, the libraries could be potentially improved with N-terminal tagging of ORFs that currently exhibit partial or no degradation of AID-tagged proteins or using multiple copies of the AID* tag to enhance protein degradation (Kubota et al., 2013; Nishimura and Kanemaki, 2014). Finally, it is noteworthy that a molecular glue for a ubiquitin ligase located in the cytosol can be used to promote degradation of the vast majority of yeast proteins, including integral membrane proteins. This suggests that targeted protein degraders such as proteolysis-targeting chimeras (PROTACs) could in principle be designed for most proteins relevant in clinical or agricultural applications based on one or a few cytosolic ubiquitin ligases (Békés et al., 2022).

## Methods

### Yeast strains and growth conditions

All yeast strains used in this work are listed in Table S6 and are derivatives of BY4741, Y8205 or Y7092. All plasmids used in this work are listed in Table S7. Yeast genome manipulations (gene tagging and gene deletion) were performed using PCR targeting and lithium acetate transformation (Janke et al., 2004; Knop et al., 1999). High-throughput N-terminal and C-terminal tagging were performed with the SWAp-Tag approach using the N-SWAT (Weill et al., 2018) and C-SWAT (Meurer et al., 2018) libraries, respectively, as detailed below. Unless stated otherwise, yeast strains were grown at 30℃ and 5-Ph-IAA (30-003-10, BioAcademia) was used at 1 μM final concentration. All strains, plasmids and libraries are available upon request.

### Construction of Halo libraries

To construct the Halo-ORF library, the donor plasmid pSD-N1-HaloTag-3myc was transformed into the donor strain yKBJ0201. The resulting strain yKBJ0203 was crossed with the N-SWAT library. In addition, to test the efficiency of tagging, the donor strain yMaM1205 was transformed with the donor plasmid pSD-N1- mNeonGreen and crossed with 13 N-SWAT strains for highly expressed proteins.

To construct the ORF-Halo library, the donor plasmid pRS41K-3myc-HaloTag was transformed into the donor strain yKBJ0200. The resulting strain yKBJ0202 was crossed with the C-SWAT library. In addition, to test the efficiency of tagging, the donor strain yKBJ0147 was transformed with the donor plasmid pRS41K- mNeonGreen and crossed with 13 N-SWAT strains for highly expressed proteins.

Crossing and subsequent tagging were performed by sequentially pinning the strains on appropriate media using a pinning robot (Rotor, Singer Instruments) in 384-colony format according to the synthetic genetic array (SGA) procedure (Baryshnikova et al., 2010a; Meurer et al., 2018; Tong et al., 2001) as follows:

- N-SWAT and C-SWAT libraries were assembled in 384-colony format on SC-Ura plates (6.7 g/l yeast nitrogen base without amino acids (BD Biosciences, 291940), 2 g/l amino synthetic complete acid mix lacking uracil (SC-Ura), 20 g/l glucose (Merck, 108337), 20 g/l agar (BD Biosciences, 214010)), whereas donor strains with the appropriate donor plasmids were pinned onto YPD+G418 plates (10 g/l yeast extract (BD Biosciences, 212750), 20 g/l peptone (BD Biosciences, 211677), 20 g/l glucose, 20 g/l agar, 200 mg/l G-418 (Biochrom, A291–25), and grown for 1-2 days at 30°C;
- mating of N-SWAT/C-SWAT and donor strains on YPD plates, 1 day at 30°C;
- selection of diploids twice on SC(MSG)-Ura + G-418 plates (1.7 g/l yeast nitrogen base without amino acids and ammonium sulfate (BD Biosciences, 233520), 1 g/l monosodium glutamic acid (MSG) (Sigma-Aldrich, G1626), 2 g/l amino acid mix SC(MSG)-Ura (glutamic acid replaced by MSG), 200 mg/l G-418, 20 g/l glucose, 20 g/l agar), 1 day at 30°C;
- sporulation on SPO plates (20 g/l potassium acetate (Sigma-Aldrich, 25059), 20g/l agar), 5-7 days at 23°C;
- selection of haploids, step 1, on SC(MSG)-Leu/Arg/Lys/Ura + canavanine/thialysine plates (50 mg/l canavanine (Sigma-Aldrich, C1625), 50 mg/l thialysine (Sigma-Aldrich, A2636)), 2 days at 30°C;
- selection of haploids, step 2, on SC(MSG)-Leu/Arg/Lys/Ura + canavanine/thialysine/G-418 plates (50 mg/l canavanine, 50 mg/l thialysine, 200 mg/l G-418), 2 days at 30°C;
- selection of haploids, step 3, on SC(MSG)-Leu/Arg/Lys/Ura + canavanine/thialysine/G-418/clonNAT plates (50 mg/l canavanine, 50 mg/l thialysine, 200 mg/l G-418, 100 mg/l clonNAT (Werner BioAgents, 5.0)), 2 days at 30°C. This step was omitted for the cross used to assess tagging efficiency with N-SWAT strains;
- selection of haploids, step 4, on SC(MSG)-Leu/Arg/Lys/Ura + canavanine/thialysine/G- 418/clonNAT/hygromycin plates (50 mg/l canavanine, 50 mg/l thialysine, 200 mg/l G-418, 100 mg/l clonNAT, 200 mg/l hygromycin (ant-hg-5, Invivogen), 2 days at 30°C. This step was omitted for the crosses used to assess tagging efficiency with N-SWAT and C-SWAT strains;
- induction of tagging twice on SC-Leu Raf/Gal plates (20 g/l galactose (Serva, 22020) and 20 g/l raffinose (Sigma-Aldrich, R0250) instead of glucose), 2 days at 30°C;
- selection against the acceptor module twice on SC-Leu + 5-FOA plates (1 g/l 5-FOA (Apollo Scientific, PC4054)), 2 days at 30°C;

Finally, the resulting strains were pinned on SC-Leu+Ade plates (300 mg/l adenine (A8626, Sigma-Aldrich)) and grown for 1 day at 30°C before proceeding with fluorescence measurements using flow cytometry and storing the libraries as glycerol stocks at -80°C.

### Construction of AID libraries

To construct the AID-v1 library, the donor plasmid pRS41K-mNeonGreen-AID*-3myc was transformed into the donor strains yEG222 (carrying the GAL1pr-OsTIR1(F74G)-natMX construct) and yMaM1205. To construct the AID-v2 library, the donor plasmid pRS41K-AID*-3myc was transformed into the donor strain yEG222. The resulting strains were crossed with the C-SWAT library. In addition, to test the efficiency of tagging, the donor strain yEG222 was transformed with the donor plasmid pRS41K-mCherry-mNeonGreen and crossed with 13 C-SWAT strains for highly expressed proteins.

Crossing and subsequent tagging were performed by sequentially pinning the strains on appropriate media in 384-colony format using the same procedure as for the Halo libraries with the following changes:

- selection of diploids twice on SC(MSG)-Ura + G-418/clonNAT plates, 1 day at 30°C;
- selection of haploids, step 1, on SC(MSG)-Leu/Arg/Lys/Ura + canavanine/ thialysine/G-418 plates, 2 days at 30°C;
- selection of haploids, step 2, on SC(MSG)-Leu/Arg/Lys/Ura + canavanine/ thialysine/G-418/clonNAT plates, 2 days at 30°C. This step was omitted for the cross used to assess tagging efficiency with the pRS41K-mCherry- mNeonGreen donor plasmid;
- induction of tagging twice on SC-Leu Raf/Gal plates (20 g/l galactose (Serva, 22020) and 20 g/l raffinose (Sigma-Aldrich, R0250) instead of glucose), 2 days at 30°C;
- selection against the acceptor module twice on SC-Leu + 5-FOA plates (1 g/l 5-FOA (Apollo Scientific, PC4054)), 2 days at 30°C;
- selection of successful tagging events with reconstitution of the hygromycin selection marker (after induction of tagging and selection against the acceptor module) twice on SC(MSG)-Leu + hygromycin/clonNAT plates (400 mg/l hygromycin, 100 mg/l clonNAT), 2 days at 30°C.

The same procedure but without clonNAT was used to construct the AID-v1 library without the GAL1pr- OsTIR1(F74G)-natMX construct.

### Competition assay

Using a pinning robot, strains from the Halo-ORF and ORF-Halo libraries were co-inoculated in 96-well plates (83.3924.005, Sarstedt) with 200 μl of SC-Leu+Ade medium per well. Each well received the two strains for a given ORF, one N-terminally tagged and carrying a cytosolic mNeonGreen marker and another C-terminally tagged and carrying a cytosolic mCherry marker. Three control wells (1 – non-fluorescent strain yMaM1205, 2 – mCherry-positive donor yKBJ0202, 3 – mNeonGreen-positive donor yKBJ0203, Table S6) served to set the gates for mCherry+ and mNG+ cells per plate. The plates were sealed with air-permeable seals (4ti-0517, 4Titude) and the co-cultures were grown overnight at 30°C to saturation.

Saturated co-cultures were diluted (5 μl of co-culture into 195 μl of SC-Leu+Ade medium) and grown for 4h at 30°C in sealed plates before single-cell fluorescence measurements with flow cytometry corresponding to day 0. After the measurements, the plates with the remaining 175 μl of co-culture per well were sealed and incubated for 20h at 30°C. The procedure starting with dilution of co-cultures was then repeated 3 times (flow cytometry analysis corresponding to days 1-3, 24h interval between measurements).

Triple competition experiments for 45 ORFs selected at random were conducted using the same procedure. Single clones of Halo-ORF and ORF-Halo strains were validated by immunoblotting of whole-cell extracts with anti-myc antibodies (mouse monoclonal, clone 9E10). Matching Halo-ORF and ORF-Halo strains were grown in each well together with a wild type strain lacking fluorescent markers (yMaM1205, Table S6).

### Flow cytometry

Strains were inoculated in 96-well plates (83.3924.005, Sarstedt) with 150-200 μl of SC-Leu+Ade medium per well and grown overnight at 30°C to saturation. Saturated cultures were diluted into fresh medium and grown at 30°C for 6 h to 6-8x10^6^ cells/ml. Assessment of tagging efficiency during construction of Halo libraries and the high-throughput competition assays were performed with strains grown in SC-Leu+Ade medium.

Single-cell fluorescence intensities were measured on a LSRFortessa SORP flow cytometer (BD Biosciences) using the BD FACSDiva v9.0.1 acquisition software, a 561 nm laser for mCherry excitation, a 600 nm long pass mirror and a 610/20 nm band pass filter for mCherry detection, a 488 nm laser for mNeonGreen excitation, a 505 nm long pass mirror and a 530/30nm band pass filter for mNeonGreen detection and a high throughput sampler with the following setting: standard mode, sample flow rate of 0.5 μl/s, sample volume of 25 μl, mixing volume of 100 μl, mixing speed of 100 μl/s, number of mixes equal to 3 and wash volume of 800 μl. 10000 cells were measured per strain. Measurements were gated for single cells with mCherry or mNeonGreen fluorescence intensities above the maximum intensity of a non-fluorescent control strain (mCherry^+^ or mNeonGreen^+^ cells). In the triple competition assay, measurements were additionally gated for cells lacking a fluorescent marker (wild type cells) (Fig. S6).

### Analysis of competition experiments

Data analysis and visualization were performed using custom scripts in R (R Core Team, 2020). Ratios of mNG^+^/mCherry^+^ cells were calculated for each well and time point from single cell fluorescence measurements obtained with flow cytometry. For each well, cell count ratios were normalized to time point zero. The relative fitness of the two strains in a well was then determined as the slope α of a linear fit to ln(normalized cell count ratio) over time, with intercept set to zero. Triple competition experiments were analyzed following the same procedure for three cell count ratios: mNG^+^/mCherry^+^ cells, mNG^+^/wild type cells and mCherry^+^/wild type cells.

Using optical density measurements at 600 nm of fifteen wells, we estimated an average doubling time of 4.5 ± 0.1h (mean ± s.d., n = 15 random ORFs) in the competition assay. The average doubling time allowed translating the relative fitness α into a difference in doubling time under the assumptions that, for a given protein, tagging of only one of the termini affects fitness and that α ≠ 0 reflects impaired fitness, not improved fitness, of one of the competing strains. Based on this and on estimates of reproducibility and technical noise in the competition assay (Fig. S1b, c), further analysis of differential fitness relied on two thresholds of abs(α) = 0.175 and 0.611 corresponding to 5% and 20% longer doubling time for one of the competing strains.

The annotation of yeast genes as essential or non-essential under standard laboratory growth conditions (Després et al., 2020; Giaever et al., 2002; Winzeler et al., 1999) and a proteome-wide localization dataset of C-terminally GFP-tagged proteins (Huh et al., 2003) were used to stratify the genome-wide distribution of relative fitness (Fig. 1c, d). Proteins assigned to multiple localization classes were counted in each class.

The relationship between differential localization and differential fitness of N- and C-terminally tagged proteins (Fig. 1e) was examined using proteome-wide localization datasets of N-terminally GFP-tagged proteins expressed from endogenous promoters and of C-terminally GFP-tagged proteins (Huh et al., 2003; Weill et al., 2018). The localization classes *missing*, *ambiguous* and *below threshold* were excluded from the analysis and only proteins with annotated localization in both datasets were considered. N- and C-terminally tagged variants assigned to multiple localization classes were considered to have the same localization if they shared at least one localization assignment (Weill et al., 2018).

The relationship between terminal localization signals and differential fitness of N- and C-terminally tagged strains was examined using a list of 998 validated or predicted localization signals obtained through literature curation or retrieved from the UniProt database (Bateman et al., 2023) (Table S2). The list consists of proteins with a glycosylphosphatidylinositol (GPI) anchor (de Groot et al., 2003; Gíslason et al., 2021), tail-anchored (TA) proteins (Beilharz et al., 2003; Burri and Lithgow, 2004; Lewis and Pelham, 2002; Schuldiner et al., 2008; Simočková et al., 2008), proteins with a cysteine-rich transmembrane module (CYSTM) (Giolito et al., 2024; Venancio and Aravind, 2010; Zvonarev et al., 2023), acylated proteins (UniProt as of 19.02.2024, search for proteins with lipidation, including N-myristoylation, palmitoylation, GPI-anchor addition and prenylation), proteins with a PTS1 peroxisomal targeting signal (Nötzel et al., 2016) (UniProt as of 24.04.2024, search for proteins with motif: microbody targeting signal, C-terminal peroxisome targeting signal (PTS1), peroxisome targeting signal (PTS1), SKL peroxisome targeting motif, peroxisomal target signal 1 (PTS1) or peroxisomal targeting signal type 1), proteins with a signal peptide (SP) or a mitochondrial targeting signal (MTS) (Somashekara and Muniyappa, 2022; Weill et al., 2018; Yofe et al., 2016), proteins with nuclear localization signals (Bordonné, 2000; Hahn et al., 2008; Lee et al., 2023; Somashekara and Muniyappa, 2022) and proteins with HDEL, KKXX or KXKXX motifs identified by pattern match or retrieved from UniProt (UniProt as of 24.04.2024, search for proteins with motif: prevents secretion from ER or di-lysine motif).

### Colony fluorescence measurements and analysis

The AID-v1 library was assembled in 1536-colony format on SC(MSG)-Leu+Ade Raf/Gal + hygromycin plates using a pinning robot (Rotor, Singer Instruments). For each ORF, six technical replicates of the *OsTIR1*^+^ strain, three technical replicates of the *OsTIR1*^-^ strain, three technical replicates of a non-fluorescent strain for background correction (AID-v2_HIS4, Table S6) and four technical replicates of an mNG^+^ reference strain (AID-v1_OsTIR1^-^_PDC1, Table S6) were arranged in a 4x4 group (Fig. 2c).

Assembled colony arrays were pinned onto SC(MSG)-Leu+Ade Raf/Gal + hygromycin + 5-Ph-IAA plates, grown for 24h at 30°C and their fluorescence was measured with a multimode microplate reader equipped with monochromators for precise selection of excitation and emission wavelengths (Spark, Tecan) and a custom temperature-controlled incubation chamber. mNG fluorescence intensities were measured with 506/5 nm excitation, 524/5 nm emission, 40 μs integration time and two detector gains for optimal detection of proteins across the full range of expression levels in the experiment. In addition, the plates were photographed with a colony imager (Phenobooth, Singer Instruments) to assess strain fitness based on colony size, as detailed in the next section.

Data analysis and visualization were performed using custom scripts in R (R Core Team, 2020). Measurements at the two detector gains were interpolated and consolidated into a single value per colony. For each ORF, measurements were normalized by the average of the reference colonies in the 4x4 group, to correct for spatial and plate effects. Sample measurements were further corrected for background fluorescence (mNG-bkg), by subtraction of the mean of the non-fluorescent colonies in the 4x4 group, or expressed in units of background fluorescence (mNG/bkg), by dividing by the mean of the non-fluorescent colonies in the group, and summarized by the mean and standard deviation of *OsTIR1*^+^ or *OsTIR1*^-^ replicates.

Background-corrected intensities of *OsTIR1*^+^ or *OsTIR1*^-^ strains were compared with a two-sided t-test and p-values were adjusted for multiple testing using the Benjamini-Hochberg method. mNeonGreen-AID*-3myc- tagged proteins were subsequently classified based on their expression levels and the extent of *OsTIR1*- dependent degradation as follows:

- not detected: mNG/bkg(*OsTIR1*^-^) ≤ 1.2;
- degraded: mNG/bkg(*OsTIR1*^-^) > 1.2, mNG/bkg(*OsTIR1*^+^) ≤ 1.2, mNG-bkg(*OsTIR1*^+^) / mNG-bkg(*OsTIR1*^-^) < 0.5 and p-value < 0.05;
- partially degraded: mNG/bkg(*OsTIR1*^-^) > 1.2, mNG/bkg(*OsTIR1*^+^*) >* 1.2, mNG-bkg(*OsTIR1*^+^) / mNG- bkg(*OsTIR1*^-^) < 0.5 and p-value < 0.05);
- not affected: mNG/bkg(*OsTIR1*^-^) > 1.2, mNG-bkg(*OsTIR1*^+^) / mNG-bkg(*OsTIR1*^-^) ≥ 0.5 or mNG/bkg(*OsTIR1*^-^) > 1.2, p-value ≥ 0.05.

Background-corrected intensities of *OsTIR1*^-^ strains were compared to measurements of absolute protein levels in molecules per cell (Lawless et al., 2016) and fluorescence measurements of a library of mNG-tagged ORFs (mNG-II library) (Meurer et al., 2018) (Fig. S2c, d). Furthermore, proteins detected in the *OsTIR1*^-^ background were split into five abundance bins of equal width based on background-corrected mNG intensities (Fig. S2f). The annotation of yeast genes as essential or non-essential under standard laboratory growth conditions (Després et al., 2020; Giaever et al., 2002; Winzeler et al., 1999), a proteome-wide localization dataset of C-terminally GFP-tagged proteins (Huh et al., 2003), a dataset of membrane protein topology (Kim et al., 2006) and annotations of essential genes as core or conditional (Peter et al., 2018; van Leeuwen et al., 2020) were used to further stratify the categorized (according to expression, extent of *OsTIR1*-dependent degradation or *OsTIR1*-dependent fitness impairment) proteins (Fig. 2e, Fig. 3d, Fig. S3c, d, Fig. S4e).

### Arrayed fitness screens and analysis

The AID-v2 library was assembled in 1536-colony format on SC(MSG)-Leu + hygromycin/clonNAT plates using a pinning robot (Rotor, Singer Instruments), with four technical replicates per strain placed next to each other in a 2x2 group. After 24h of growth, the colony arrays were pinned on different media with 1 μM 5-Ph-IAA and genotoxic agents (GAs): control (YP Raf/Gal + hygromycin/clonNat), 5-Ph-IAA (YP Raf/Gal + hygromycin/clonNat/5-Ph-IAA), GA (YP Raf/Gal + hygromycin/clonNat/GA) and 5-Ph-IAA + GA (YP Raf/Gal + hygromycin/clonNat/5-Ph-IAA/GA) plates, where GA stands for camptothecin (CPT, 250 μM; 7689-03-4, Sigma-Aldrich), hydroxyurea (HU, 150 mM; 127-07-1, Sigma-Aldrich) or methyl methanesulfonate (MMS, 0.02% v/v; 66-27-3, Sigma-Aldrich).

Colony arrays were grown at 30°C and photographed after 24h and 48h with a colony imager (Phenobooth, Singer Instruments). Subsequent data analysis and visualization were performed using custom scripts in R (R Core Team, 2020). The gitter package was used to segment the photographs and determine colony sizes (Wagih and Parts, 2014). The SGA tools package was used to correct for spatial and plate effects (Wagih et al., 2013): four outer rows and columns of dummy colonies on each plate were removed; spatial correction was performed without the row and column corrections; corrected measurements were normalized by the median of each plate (median calculated using only colonies between the 40th and 80th percentiles due to the high number of strains with impaired fitness). For each condition, the measurements were then normalized by the median of that condition (median of the data between the 40th and 80th percentiles) and summarized by the mean and standard deviation for each strain.

A similar procedure was used to determine the colony sizes in the AID-v1 library grown according to Fig. 2c. Next, the impact of the *GAL1-OsTIR1(F74G)* construct was estimated at 0.93 as the colony size of an *OsTIR1*^+^ strain relative to the corresponding *OsTIR1*^-^ strain, averaged across non-essential ORFs (Fig. S3a). Colony size measurements of *OsTIR1*^+^ strains were then corrected for the fitness impact of OsTir1 expression using the multiplicative model (Baryshnikova et al., 2010b), whereby the fitness impacts of OsTir1 expression and degradation of the AID-tagged protein are independent: size(*OsTIR1*^+^, corrected) = size(*OsTIR1*^+^) / 0.93.

To identify strains with impaired growth upon degradation of the AID-tagged protein, corrected colony sizes of *OsTIR1*^+^ and normalized colony sizes of *OsTIR1*^-^ strains in the AID-v1 library or normalized colony sizes of AID-v2 strains on control and 5-Ph-IAA plates were compared with a two-sided t-test and p-values were adjusted for multiple testing using the Benjamini-Hochberg method. Strains with impaired fitness were defined as follows:

- AID-v1 library: size(*OsTIR1*^+^)/size(*OsTIR1*^-^) < 0.8 and p-value < 0.05 (Fig. 3a);
- AID-v2 library: size(5-Ph-IAA)/size(control) < 0.8 and p-value < 0.05 (Fig. S4b).

Chemical-genetic interaction (CGI) scores were calculated for each ORF using the normalized median-centered colony sizes and the multiplicative model (Baryshnikova et al., 2010b) as follows:

CGI score = size(5-Ph-IAA, GA) – size(5-Ph-IAA) x size(GA).

Finally, colony sizes in the double perturbation condition (5-Ph-IAA, GA) were compared against the product of sizes from the single perturbation (5-Ph-IAA and GA) with a two-sided t-test and p-values were adjusted for multiple testing using the Benjamini-Hochberg method. Significant negative chemical-genetic interactions were defined as follows: size(5-Ph-IAA, GA) ≤ 0.9, CGI score ≤ -0.2 and p-value < 0.05. The identified genes were mapped to gene ontology (GO) terms using the GO slim mapper in the Saccharomyces genome database with GO version 2024-01 (Wong et al., 2023) and grouped into broader functional categories (Fig. 4d, Fig. S5c) as follows:

- transcription GO terms: transcription by RNA polymerase I, transcription by RNA polymerase II, mRNA processing, rRNA processing and RNA catabolic process;
- chromatin organization GO terms: chromatin organization;
- DNA damage response GO terms: DNA repair, DNA recombination and DNA damage response;
- cell cycle GO terms: meiotic cell cycle, mitotic cell cycle, regulation of cell cycle, DNA replication, sporulation and chromosome segregation;
- metabolism GO terms: amino acid metabolic process, vitamin metabolic process, generation of precursor metabolites and energy, carbohydrate metabolic process and nucleobase-containing small molecule metabolic process;
- proteostasis GO terms: protein glycosylation, protein folding, protein modification by small protein conjugation or removal, and proteolysis involved in protein catabolic process;
- intracellular trafficking and lipid metabolism GO terms: endosomal transport, transmembrane transport, vacuole organization, endocytosis, golgi vesicle transport, membrane fusion, vesicle organization, lipid metabolic process and protein targeting.

### Yeast spot assays

Strains were grown overnight in YPD medium at 30°C. Ten-fold serial dilutions, starting with 0.5 OD600nm, were spotted onto agar plates using a replica plater (R2383, Sigma-Aldrich) and incubated at 30°C. The plates were photographed (ChemiDoc Touch imaging system, Bio-Rad) after 2-4 days of incubation.

### Time courses of inducible protein degradation

Strains were grown overnight in YP Raf medium at 30°C, diluted to 0.2 OD600nm and incubated for 3h, followed by addition of galactose to 2% (w/v) to induce expression of *OsTIR1*. Time courses were initiated 90 min later with addition of 5-Ph-IAA to 1 μM final concentration and 1.0 OD600nm samples were collected at 0, 10, 30 and 60 min time points.

Whole cell extracts were prepared by alkaline lysis followed by precipitation with trichloroacetic acid (Knop et al., 1999) and separated by SDS-PAGE, followed by immunoblotting against the myc tag and against the loading control Pgk1 with mouse anti-myc (1:1000 dilution, clone 9E10, IMB Protein Production core facility), mouse anti-Pgk1 (1:10000 dilution, 459250, ThermoFisher Scientific) and goat HRP-conjugated anti-mouse (1:3000 dilution, 1705047, Bio-rad) antibodies. Membranes were developed using a chemiluminescent horseradish peroxidase substrate (SuperSignal West Pico PLUS, 34580, ThermoFisher Scientific) and imaged using a ChemiDoc Touch imaging system (Bio-Rad).

### Cell cycle profiling

Overnight cultures in YP Raf medium were diluted to 0.2 OD600nm and incubated at 30°C. After 3h, galactose was added to 2% (w/v) to induce expression of *OsTIR1*. Time courses were initiated 90 min later with the addition of HU to 150 mM final concentration and 1 ml samples were collected every 15 min for 2h. At the 2h time point, cultures were pelleted by centrifugation, washed three times with pre-warmed YP Raf/Gal and released into YP Raf/Gal medium at 25°C. To follow the release from HU, samples were collected every 15 min for 90 min.

At each time point, cells were harvested by centrifugation and fixed in 70% ethanol overnight at 4°C. Fixed cells were pelleted by centrifugation at 17000 g for 5 min, washed once with 800 μl of 50 mM sodium citrate pH 7.4 buffer, resuspended in 500 μl of 50 mM sodium citrate pH 7.4 buffer, treated with 0.25 mg/ml RNase A (10753721, ThermoFisher Scientific) for 3h at 37°C, followed by treatment with 1 mg/ml Proteinase K (Genaxxon, M3037.0005) for 2h at 50°C. Treated samples were stained by addition of SYTOX Green (1076273, ThermoFisher Scientific) to 4 μM final concentration. Cell cycle profiles were measured with a flow cytometer (LSRFortessa, BD Biosciences with BD FACSDiva software v9.7), with 10000 recorded events per sample. Data analysis was performed with FlowJo v10.10 (BD Biosciences).

## Data Availability

The results of the genome-wide competition screen, characterization of the AID-v1 and AID-v2 library, and of the genome-wide screens for DNA damage response factors are provided as supplementary tables.

## Code Availability

The pipelines for downstream analysis and data visualization, together with the necessary input data, are available at https://github.com/khmelinskii-lab/AID_libraries.

## Acknowledgements

We thank Helle Ulrich for reagents. We thank the IMB Flow Cytometry core facility for the use of their instruments and Stefanie Möckel for help with the experimental setup of the competition screen and initial data analysis, the IMB Protein Production core facility for the mouse anti-myc antibody. Funding of the German Research Foundation supported the BD LSRFortessa SORP (P#210253511). The IMB Media Lab and the IMB Protein Production facility are gratefully acknowledged for support with growth media and reagents. This work was supported by the European Research Council (ERC-2017-STG#759427 to A.K.).

## Author Contributions Statement

Conceptualization: BL, AK; experimental investigation: EG, KAJN; data analysis: EG, KAJN, JJF, SS; writing-original draft: AK; writing-review and editing: AK, BL, EG, KAJN; supervision: BL, AK.; funding acquisition: BL, AK.

## Supplementary Tables

**Supplementary Table 1.** Results of the genome-wide competition screen.

**Supplementary Table 2.** Terminal localization signals in the yeast proteome.

**Supplementary Table 3.** Characterization of the AID-v1 library.

**Supplementary Table 4.** Results of the genome-wide DDR screens.

**Supplementary Table 5.** Gene set enrichment analysis of resistance factors unique to each genotoxic agent

**Supplementary Table 6.** Yeast strains.

| Strain | Genotype | Source |
| --- | --- | --- |
| BY4741 | MATa his3Δ1 leu2Δ0 met15Δ0 ura3Δ0 | (Brachmann et al., 1998) |
| Y8205 | MATalpha can1Δ::STE2pr-SpHIS5 lyp1Δ::STE3pr-LEU2 his3Δ1 leu2Δ0 ura3Δ0 | (Tong and Boone, 2007) |
| Y7092 | MATalpha lyp1Δ his3Δ1 leu2Δ0 ura3Δ0 met15Δ0 can1Δ::STE2pr-SpHIS5 | (Tong and Boone, 2007) |
| yMaM1205 | MATalpha lyp1Δ his3Δ1 leu2Δ0 ura3Δ0 met15Δ0 can1Δ::STE3pr-LEU2-GAL1pr-NLS-I-SCEI | (Meurer et al., 2018) |
| yKBJ0147 | yMaM1205 leu2Δ::natNT2-TEF1pr-mCherry-CYC1term | this study |
| yKBJ0148 | yMaM1205 leu2Δ::natNT2-TEF1pr-mNeonGreen-CYC1term | this study |
| yKBJ0200 | yMaM1205 leu2Δ::natNT2-TEF1pr-mCherry-CYC1term pdr5Δ::hphNT1 | this study |
| yKBJ0201 | yMaM1205 leu2Δ::natNT2-TEF1pr-mNeonGreen-CYC1term pdr5Δ::hphNT1 | this study |
| yKBJ0202 | yMaM1205 leu2Δ::natNT2-TEF1pr-mCherry-CYC1term pdr5Δ::hphNT1 pRS41K-3myc-HaloTag | this study |
| yKBJ0203 | yMaM1205 leu2Δ::natNT2-TEF1pr-mNeonGreen-CYC1term pdr5Δ::hphNT1 pSD-N1-HaloTag-3myc | this study |
| yEG55 | yMaM1205 lyp1Δ::GPDpr-OsTIR1(F74G)-natMX | this study |
| yEG222 | yMaM1205 lyp1Δ::GAL1pr-OsTIR1(F74G)-natMX | this study |
| yEG224 | yMaM1205 lyp1Δ::CYC1pr-OsTIR1(F74G)-natMX | this study |
| yEG292 | yMaM1205 lyp1Δ::ADH1pr-OsTIR1(F74G)-natMX | this study |
| AID-v1 library (OsTIR1<sup>-</sup> set) | MATa lyp1Δ his3Δ1 leu2Δ0 ura3Δ0 met15Δ0 can1Δ::STE3pr-LEU2-GAL1pr-NLS-I-SCEI ORF-mNeonGreen-AID*-3myc-hph | this study |
| AID-v1 library (OsTIR1<sup>+</sup> set) | MATa his3Δ1 leu2Δ0 ura3Δ0 met15Δ0 can1Δ::STE3pr-LEU2-GAL1pr-NLS-I-SCEI lyp1Δ::GAL1pr-OsTIR1(F74G)-natMX ORF-mNeonGreen-AID*-3myc-hph | this study |
| AID-v2 library | MATa his3Δ1 leu2Δ0 ura3Δ0 met15Δ0 can1Δ::STE3pr-LEU2-GAL1pr-NLS-I-SCEI lyp1Δ::GAL1pr-OsTIR1(F74G)-natMX ORF-AID*-3myc-hph | this study |
| AID-v1_OsTIR1<sup>-</sup>_PDC1 | MATa lyp1Δ his3Δ1 leu2Δ0 ura3Δ0 met15Δ0 can1Δ::STE3pr-LEU2-GAL1pr-NLS-I-SCEI PDC1-mNeonGreen-AID*-3myc-hph | this study |
| AID-v2_HIS4 | MATa his3Δ1 leu2Δ0 ura3Δ0 met15Δ0 can1Δ::STE3pr-LEU2-GAL1pr-NLS-I-SCEI lyp1Δ::GAL1pr-OsTIR1(F74G)-natMX HIS4-AID*-3myc-hph | this study |

**Supplementary Table 7.** Plasmids.s

| Plasmid | Description | Source |
| --- | --- | --- |
| pSD-N1 | CEN ARS kanMX (empty type I N-SWAT donor) | (Yofe et al., 2016) |
| pMaM482 | pRS41K CEN ARS kanMX (empty type I C-SWAT donor) | (Meurer et al., 2018) |
| pEI098 | pSD-N1-mNeonGreen (N-SWAT type I donor, tagging with mNeonGreen) | this study |
| pYD10 | pRS41K-mNeonGreen (C-SWAT type I donor, tagging with mNeonGreen) | (Meurer et al., 2018) |
| pKBJ050 | pSD-N1-HaloTag-3myc (N-SWAT type I donor, tagging with HaloTag-3myc) | this study |
| pKBJ051 | pRS41K-3myc-HaloTag (C-SWAT type I donor, tagging with 3myc-HaloTag) | this study |
| pBL1099 | pRS41K -AID*-3myc (C-SWAT type II donor, tagging with AID*-3myc) | this study |
| pBL1101 | pRS41K-mNeonGreen-AID*-3myc (C-SWAT type II donor, tagging with mNeonGreen-AID*-3myc) | this study |

**Figure S1.**
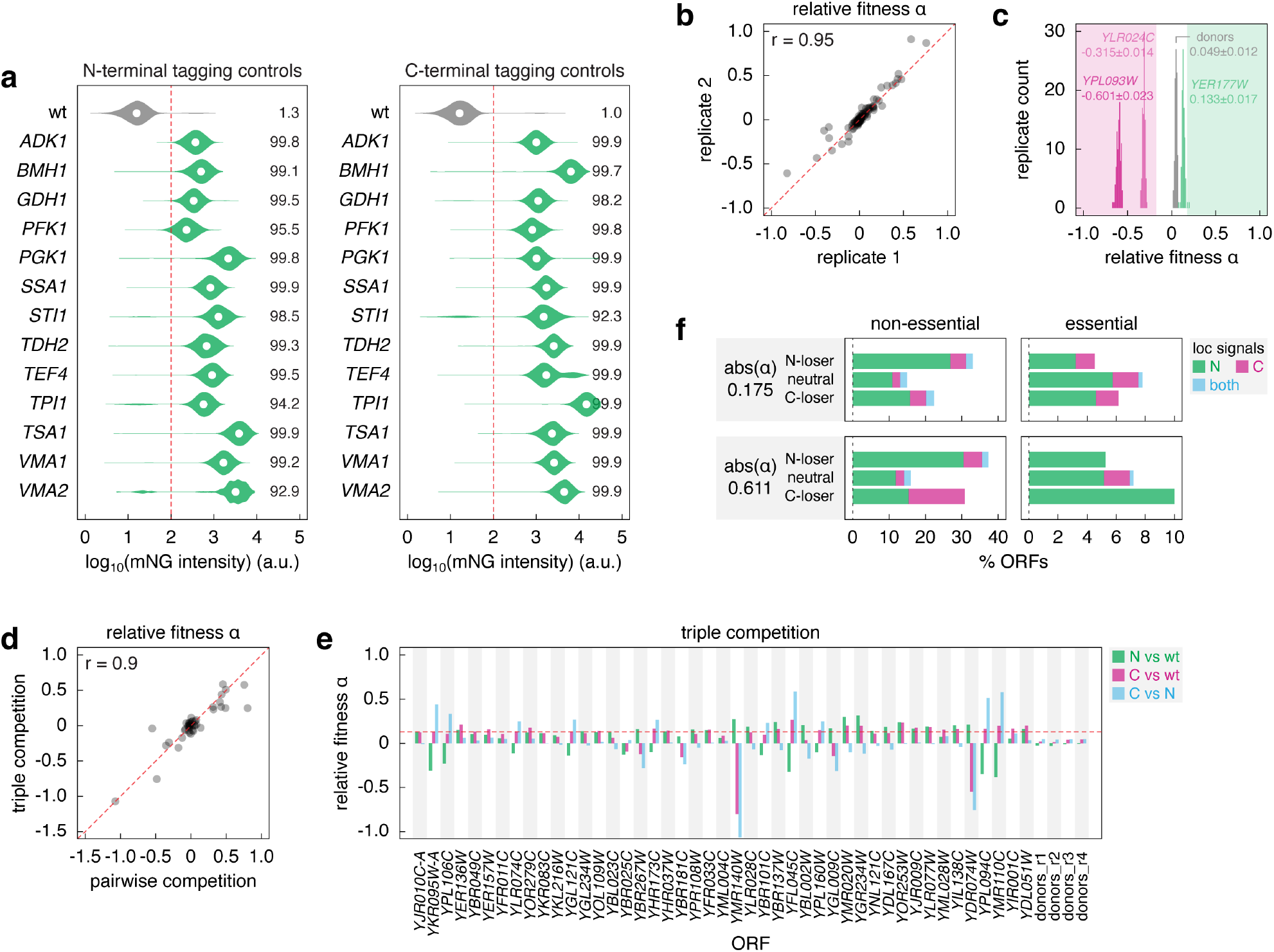
Fitness impact of N- vs C-terminal tagging. **a** – Evaluation of the SWAT tagging efficiency during construction of the Halo-ORF and ORF-Halo libraries using donors with mNeonGreen (mNG) as the tag. Tagging was performed with 13 N-SWAT (left) or C-SWAT (right) strains for the indicated ORFs. Distributions of single-cell mNeonGreen fluorescence intensities measured with flow cytometry (10000 cells per strain). The percentage of cells with fluorescence above background (fluorescence of a wild type strain, red dashed line) is indicated. **b** – Reproducibility of relative fitness estimation with the competition assay. Relative fitness of two replicates for 92 ORFs, determined as in Fig. 1a. r, pearson correlation coefficient. **c** – Evaluation of technical noise in the competition assay. Distributions of relative fitness determined as in Fig. 1a for the indicated ORFs and the pair of SWAT donor strains (no tagged ORF), 92 technical replicates per strain pair. Mean ± s.d. relative fitness is indicated. **d, e** – Triple competition of 45 randomly selected pairs of Halo-ORF and ORF-Halo strains against a wild type strain lacking fluorescent markers. Relative fitness of Halo-tagged strains determined according to Fig. 1a and in the triple competition assay (**d**, Methods). Pairwise relative fitness of the three strains for each ORF in the triple competition assay (**e**, Methods). Median relative fitness (0.13) of N- and C-terminal variants against wild type (red dashed line, **e**). **f** – Proteins with localization signals at the N-terminus, C-terminus or both protein termini, stratified by gene essentiality and differential fitness according to Fig. 1b. Analysis based on 997 validated or predicted localization signals: 611, 188 and 99 proteins with signals at the N-terminus, the C-terminus or at both termini, respectively (Methods, Table S2). N-loser, neutral and C-loser ORFs defined at two differential fitness thresholds.

**Figure S2.**
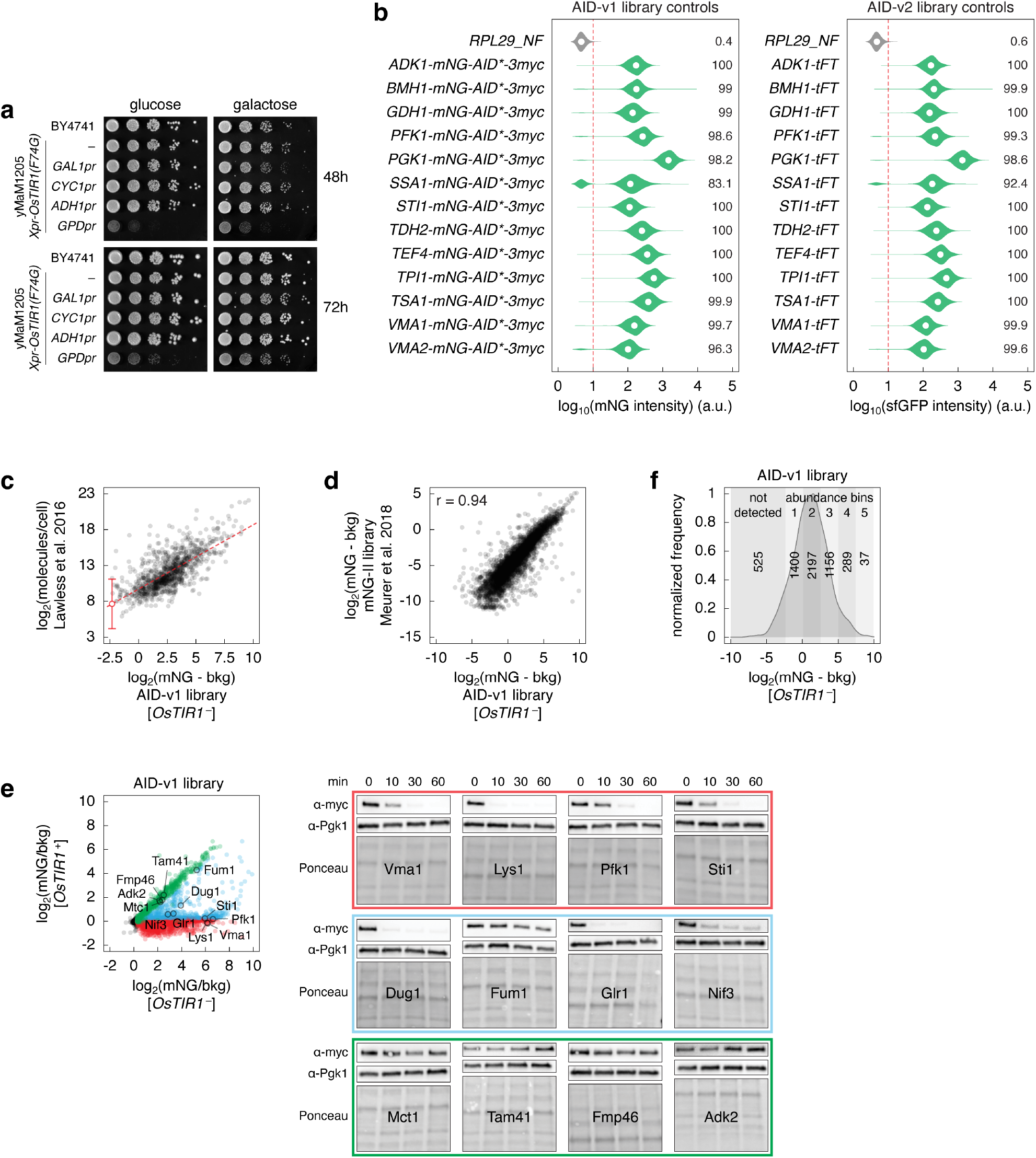
Construction and characterization of genome-wide AID libraries. **a** – Fitness impact of *OsTIR1* expression from different promoters. Ten-fold serial dilutions of yeast strains on the indicated media without 5-Ph-IAA. **b** – Evaluation of the SWAT tagging efficiency during construction of the AID libraries. For the AID-v2 library, control tagging was performed with 13 C-SWAT strains for the indicated ORFs and a donor with the mCherry- sfGFP timer (tFT) as a tag. Distributions of single-cell mNeonGreen fluorescence intensities measured with flow cytometry (10000 cells per strain). The percentage of cells with fluorescence above background (fluorescence of a wild type strain, red dashed line) is indicated. **c** – Correlation between absolute protein abundance (Lawless et al., 2016) and relative protein expression levels measured with the AID-v1 library lacking *OsTIR1*. Fluorescence intensities of colonies were corrected for background fluorescence. Linear fit (red dashed line) and estimate of the detection limit of the colony assay (200 molecules/cell) with the 95% confidence interval (18 to 2187 molecules/cell). **d** – Comparison of relative protein expression levels measured with the AID-v1 library lacking *OsTIR1* and with the mNG-II library (ORF-mNG strains) (Meurer et al., 2018). Fluorescence intensities of colonies were corrected for background fluorescence. **e** – Degradation of AID-tagged proteins after addition of 5-Ph-IAA. Whole-cell extracts of *OsTIR1*^+^ strains for the indicated proteins (AID-v1 library, left) were separated by SDS-PAGE, followed by immunoblotting with antibodies against the myc tag and Pgk1 as the loading control (right). **f** – Distribution of background-corrected mNG fluorescence intensities in the AID-v1 library, *OsTIR1*^-^ strains, determined according to Fig. 2c. Number of proteins in the not detected group (mNG/bkg(*OsTIR1*^-^) ≤ 1.2) and in five abundance bins used in Fig. 2e, Fig. 3d, Fig. S3b and Fig. S4e is indicated.

**Figure S3.**
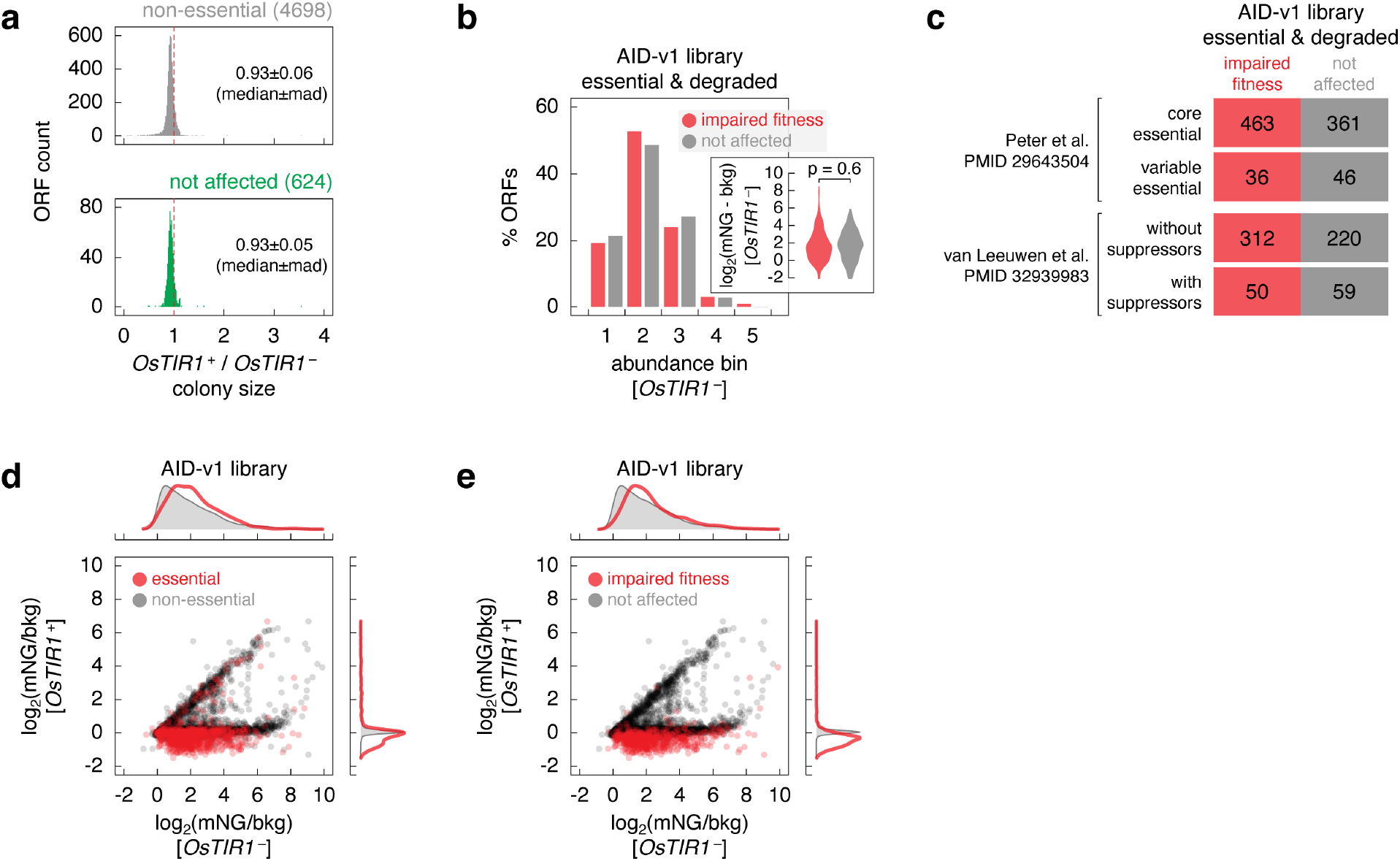
Characterization of the AID-v1 library. **a** – Fitness impact of the *GAL1pr-OsTIR1(F74G)* expression construct. For each ORF, the fitness impact was estimated as the colony size of the *OsTIR1*^+^ strain relative to the *OsTIR1*^-^ strain in the AID-v1 library (*OsTIR1*^+^/*OsTIR1*^-^), determined according to Fig. 2c. Distributions of relative colony sizes for two sets of ORFs (4698 non-essential ORFs or 624 not affected proteins in Fig. 2d). Median ± mad (median absolute deviation) relative fitness is indicated. **b** – Expression levels of essential and degraded proteins in the AID-v1 library (Fig. 2c), according to fitness of *OsTIR1*^+^ strains. Protein abundance (mNG signal corrected for background fluorescence, mNG-bkg) in the *OsTIR1*^-^ background; abundance bins defined in Fig. S2f; p-value in a Mann-Whitney U test. **c** – Number of *OsTIR1*^+^ strains in the AID-v1 library with essential and degraded proteins, stratified by essential gene type: core or variable according to (Peter et al., 2018), with or without suppressors according to (van Leeuwen et al., 2020). **d, e** – Protein levels (mNG signal in units of background fluorescence, mNG/bkg) in the AID-v1 library determined according to Fig. 2c. Essential and non-essential ORFs (**d**) and strains with normal or impaired fitness based on colony size measurements (**e**) are marked. Top and right, normalized frequency distributions of protein levels in each category.

**Figure S4.**
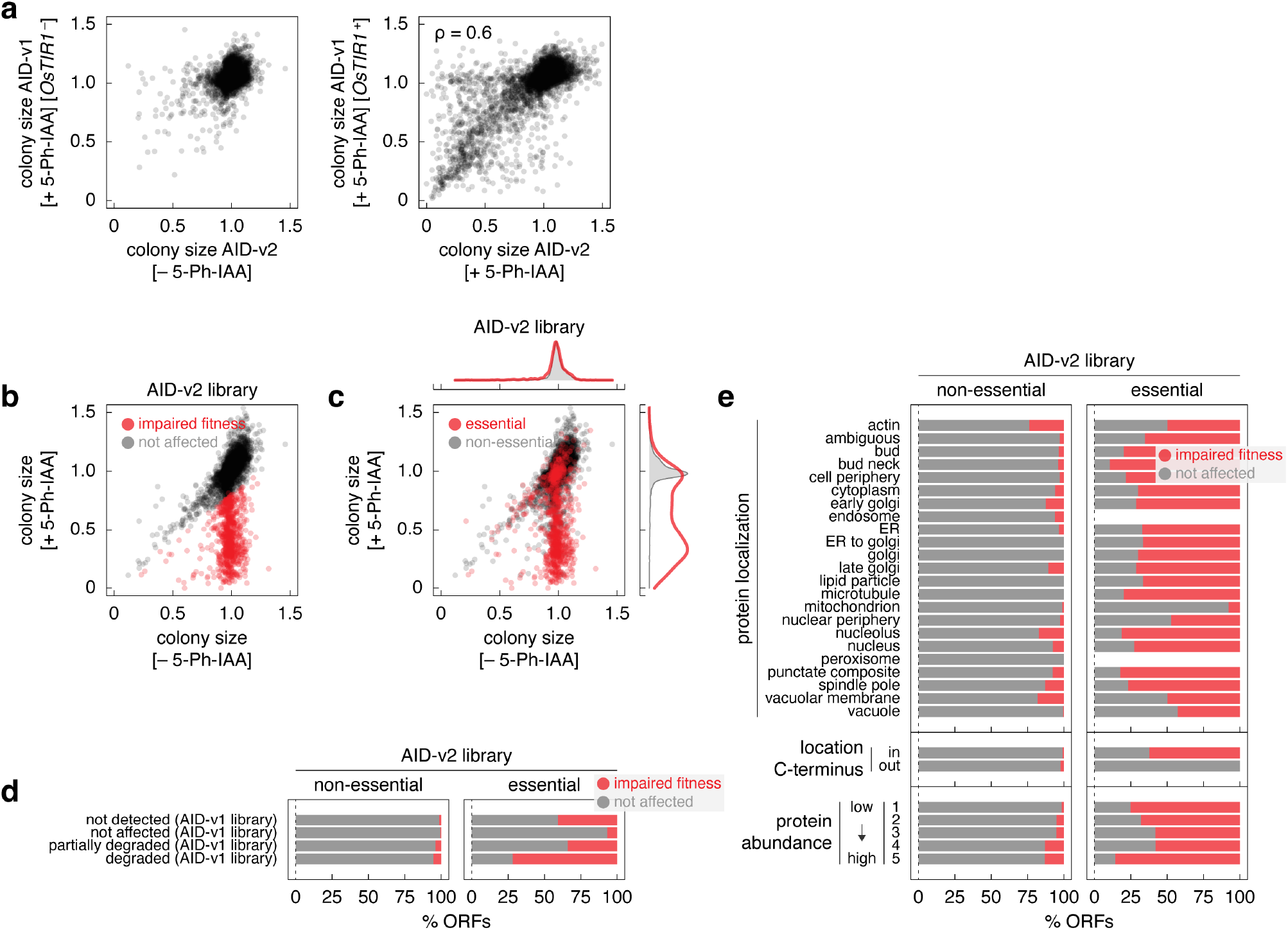
Characterization of the AID-v2 library. **a** – Comparison of normalized colony sizes in the AID-v1 and AID-v2 libraries grown according to Fig. 2c and Fig. 4a, respectively. ρ, spearman correlation coefficient. **b, c** – Normalized colony sizes in the AID-v2 library grown according to Fig. 4a on galactose medium with or without 1 μM 5-Ph-IAA. **c**, essential and non-essential ORFs are marked; top and right, normalized frequency distributions of normalized colony sizes in each category. **d, e** – Frequency of 5-Ph-IAA-dependent fitness defects in the AID-v2 library according to the extent of *OsTIR1*-dependent degradation phenotypes in the AID-v1 library (**d**) or according to protein localization, location of the C-terminus (cytosol – in, lumen of organelles or extracellular – out) or protein abundance estimates from the AID1-v1 library (**e**).

**Figure S5.**
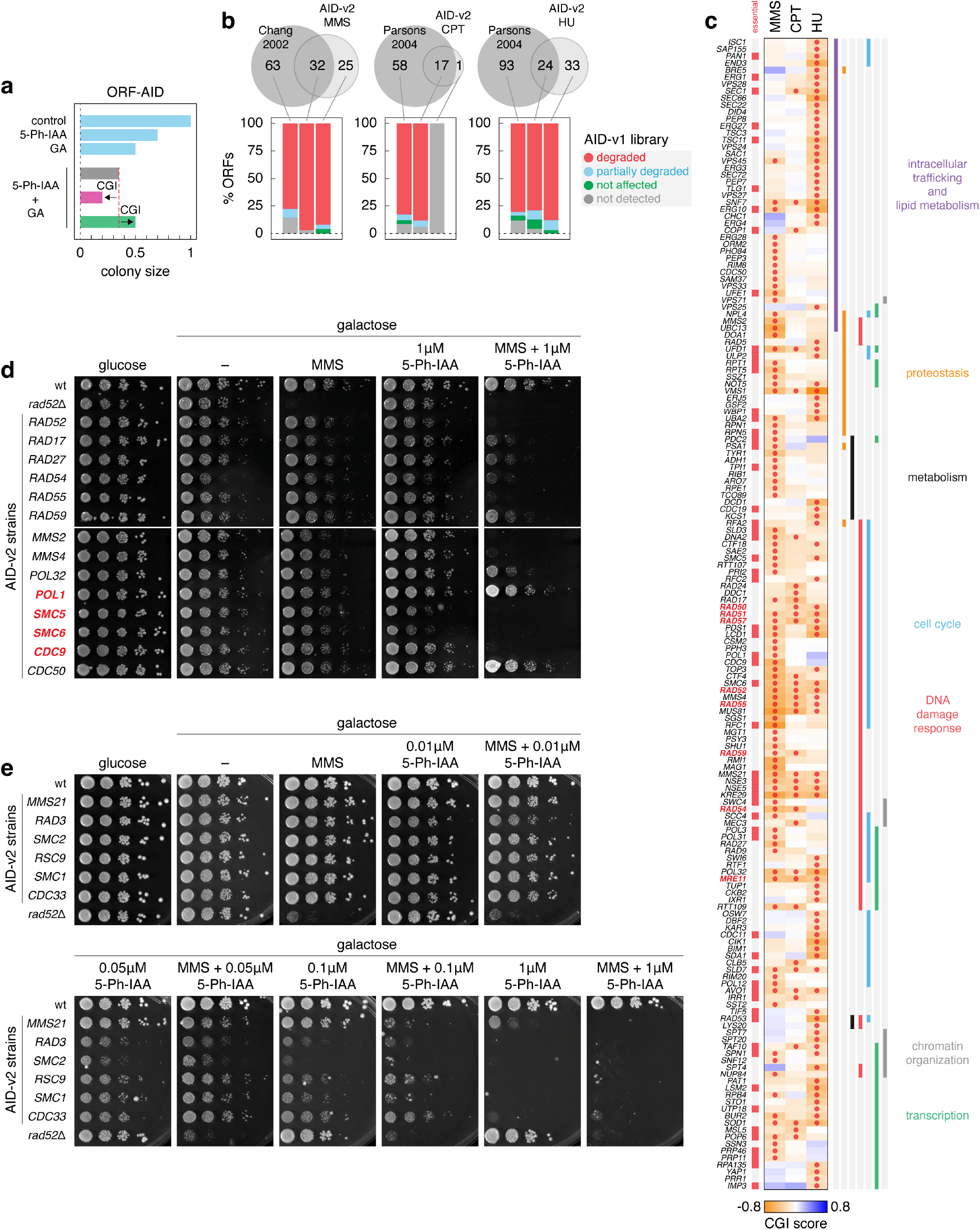
Genome-wide screens for DNA damage response factors. **a** – Definition of chemical-genetic interaction (CGI). Fitness of an ORF-AID strain (normalized colony size, Methods) measured under four conditions: control (no perturbation), in the presence of 5-Ph-IAA (to induce degradation of the AID-tagged protein), GA (genotoxic agent) or both. Grey bar, expected fitness in the double perturbation condition (product of the fitness from the single perturbation conditions) in the absence of a CGI. Red and green bars, examples of negative and positive CGIs. **b** – Top, overlap between chemical-genetic interactors identified with the AID-v2 library and previously identified resistance genes for MMS (Chang et al., 2002), CPT or HU (Parsons et al., 2003). Essential factors identified with the AID-v2 library were excluded from the comparison. Bottom, frequency of *OsTIR1*-dependent degradation phenotypes in the AID-v1 library for the indicated sets of genes. **c** – Heatmap of negative chemical-genetic interactors with MMS, CPT or HU, as in Fig. 4d. Gene names, essential genes and significant interactors (red circles) with each genotoxic agent are indicated. **d, e** – Ten-fold serial dilutions of the indicated strains from the AID-v2 library, a wild type (wt) and a *RAD52* knockout on the indicated media. Essential genes are marked in red (**d**).

**Figure S6.**
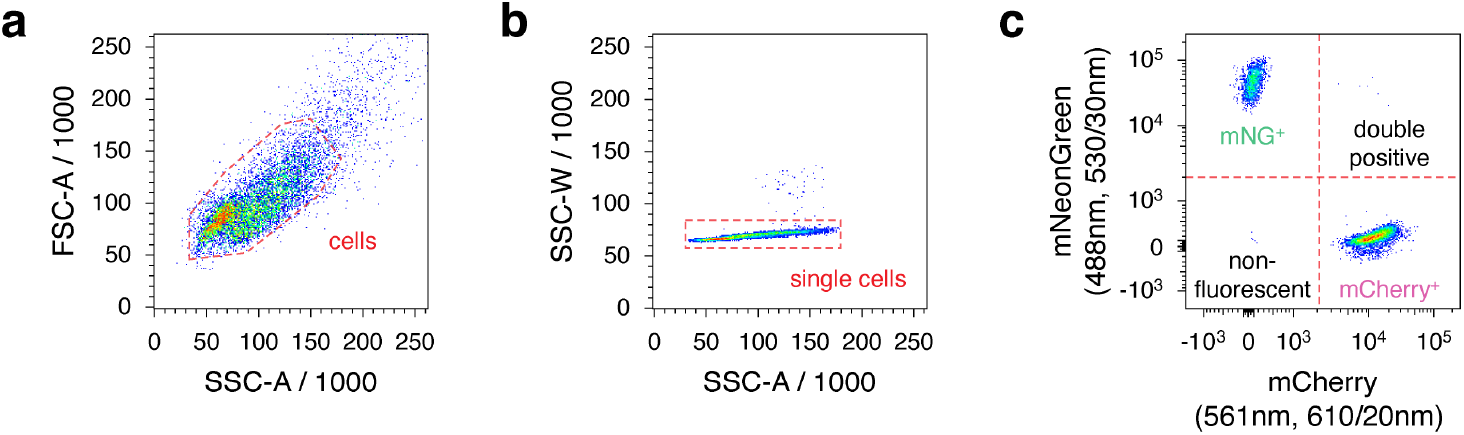
Representative gating strategy used in flow cytometry experiments. **a** – Measurements were gated to identify cells based on size and granularity using forward and side scatter areas. **b** – Cells were subsequently gated for single cells using side scatter area and width. **c** – Finally, single cells were gated into four populations based on their mNeonGreen or mCherry fluorescence intensities compared to the maximum intensities of a non-fluorescent control population.

